# Testing heterochrony: Connecting skull shape ontogeny and evolution of feeding adaptations in baleen whales

**DOI:** 10.1101/2022.11.29.518348

**Authors:** Agnese Lanzetti, Roberto Portela Miguez, Vincent Fernandez, Anjali Goswami

## Abstract

Ontogeny plays a key role in the evolution of organisms, as changes during the complex processes of development can allow for new traits to arise. Identifying changes in ontogenetic allometry – the relationship between skull shape and size during growth – can reveal the processes underlying major evolutionary transformations. Baleen whales (Mysticeti, Cetacea) underwent major morphological changes in transitioning from their ancestral raptorial feeding mode to the three specialised filter feeding modes observed in extant taxa. Heterochronic processes have been implicated in the evolution of these feeding modes, and their associated specialised cranial morphologies, but their role has never been tested with quantitative data. Here, we quantified skull shapes ontogeny and reconstructed ancestral allometric trajectories using 3D geometric morphometrics and phylogenetic comparative methods on sample representing modern mysticetes diversity. Our results demonstrate that Mysticeti, while having a common developmental trajectory, present distinct cranial shapes from early in their ontogeny corresponding to their different feeding ecologies. Size is the main driver of shape disparity across mysticetes. Disparate heterochronic processes are evident in the evolution of the group: skim feeders present accelerated growth relative to the ancestral nodes, while Balaenopteridae have overall slower growth, or paedomorphosis. The grey whale is the only taxon with a relatively faster rate of growth in this group, which might be connected to its unique benthic feeding strategy. Reconstructed ancestral allometries and related skull shapes indicate that extinct taxa used less specialized filter feeding modes, a finding broadly in line with the available fossil evidence.

## 1 Introduction

The intrinsic relationship between ontogeny and phylogeny has long been explored, with Darwin himself viewing changes in development as an argument in favour of his theory of descent with modification (Haeckel, 1866; Gould, 1977). Possibly the most important concept to bridge evolution and development is heterochrony, broadly defined as changes in the relative timing of appearance or growth rate of characters between the ancestor and its descendants (Alberch et al., 1979; McNamara, 1986; Klingenberg, 1998). Heterochronic processes can act on multiple scales, from gene expression to the development of entire phenotypic structures (e.g., skull) (Smith, 2003; Olsson et al., 2017). These changes can be divided into two broad categories based on their effect on the descendant morphology: paedomorphosis, in which the adult organism retains juvenile characters of the ancestor, and peramorphosis, in which the juvenile of the descendant presents traits that characterized the adult ancestor (Alberch et al., 1979; McNamara, 1986; Klingenberg, 1998). Quantifying heterochrony in the evolution of a group allows one to assess the role of development in the rise of new traits and adaptations in both living and extinct organisms and clarify phylogenetic relationships based on morphological traits (Wiens et al., 2005; Thewissen et al., 2012). A well-studied lineage that shows examples of both types of heterochrony acting at different levels is birds (Aves). The skull of modern birds appears to possess an overall paedomorphic shape, while their postcranial skeleton presents peramorphic features, both shifts relative to extinct non-avian dinosaurs (Bhullar, 2012; Bhullar et al., 2016). These heterochronic changes are hypothesized to be related to locomotory and dietary transformations that occurred in the evolution of this clade (Bhullar et al., 2016).

An important developmental concept that is connected with heterochronic processes is allometry, defined as the relationship between changes in size and shape of a structure during the growth of an organism (Gould, 1977; McNamara, 1986; Klingenberg, 1998). Allometric trajectories are a powerful tool for understanding heterochrony as they can be recorded during the growth of a single organism or across species without data on raw or relative developmental timing (McKinney, 1988). A clear instance of evolutionary allometry is the gigantism or extreme increase in body size in sauropods (herbivorous non-avian dinosaurs), achieved through peramorphosis and probably connected to shifts in global climate and resource availability (Sander et al., 2011; Clauss et al., 2013). On the other hand, a typical case of paedomorphosis in evolutionary allometry is the dwarfism or reduction of body size, as well as brain size, in many mammalian insular species, such as hippos, elephants and humans, which was likely caused by the lack of predators and decrease in diet quality (Weston and Lister, 2009; Herridge and Lister, 2012). Identifying and quantifying patterns of growth and evolutionary allometry and how they act on different parts of the organism’s phenotypes is a key part of studying the role of heterochrony in the evolution of a lineage (McNamara, 1986; McKinney, 1988; Klingenberg, 1998).

In Cetacea, including the extant toothed (Odontoceti) and baleen whales (Mysticeti), as well as the extinct archaeocetes, several recent studies have investigated different aspects of the contribution of ontogeny to the evolution of this unique mammalian clade. The loss of functional hind limbs in cetaceans is a key adaptation to life in the water, allowing more efficient swimming modes (Thewissen and Fish, 1997), and has been linked to changes in timing and duration of regulatory gene expression early in ontogeny (Thewissen et al., 2006). Alterations in location and timing of gene expression have also been implicated in the evolution of homodont dentition in Odontoceti and in the growth of baleen during the prenatal development of modern Mysticeti (Armfield et al., 2013; Thewissen et al., 2017). Heterochrony on a broader, phenotypic level has also been found to have had a strong impact on the phylogeny and diversification of Odontoceti, especially in porpoises and at least one dolphin genus (*Cephalorhynchus*) (Galatius and Kinze, 2003; Galatius, 2010; Galatius et al., 2011), including by influencing their level of skull asymmetry and related hearing adaptations (Lanzetti et al., 2022).

In Mysticeti, several studies have focused on postnatal growth and evolutionary allometry of the skull and its various specialised structures (e.g., Goldbogen et al., 2010; Nakamura et al., 2012; Pyenson et al., 2013; Franci and Berta, 2018; Kahane-Rapport and Goldbogen, 2018; Werth et al., 2018). For example, it has been shown that filtration area, measured as surface area of baleen, is positively correlated with body size in all living mysticetes (Werth et al., 2018). Different families with diverse filter feeding styles have unique allometric scaling of skull and body size as they are constrained by the shape of their rostrum and of the baleen (Kahane-Rapport and Goldbogen, 2018).

Balaenidae (*Balaena mysticetus* and *Eubalaena* spp.), the group that includes bowhead and right whales, use a skim feeding strategy, swimming slowly through the water with their mouth open and capture plankton, copepods and amphipods mostly, using their long baleen plates (Berta et al., 2016). Thanks to their arched upper jaw that allows them to accommodate the long baleen (Bouetel, 2005), they have a larger buccal cavity relative to body size compared to other living mysticetes. Rorqual whales (Balaenopteridae) employ an engulfment or lunge feeding strategy: they feed in gulps, by scooping a large amount of prey – predominantly schooling fish – in their mouth either by swimming horizontally (engulfment) or by lunging out of the water (lunge). They have a broad and flat rostrum, built to resist the impact of the water during feeding, and blunt baleen plates (Berta et al., 2016).

Their head size, mandibular length and buccal capacity have been shown to increase allometrically with body size, with larger species having a greater head-to-body size ratio than smaller ones (Goldbogen et al., 2010; Pyenson et al., 2012; Kahane-Rapport and Goldbogen, 2018). This relationship likely has driven the evolution of this group, with taxa progressively modifying their body proportions over time to best adapt to different ecological niches. For example, smaller species like the minke whales (*B. acutorostrata* and *B. bonaerensis*) have higher manoeuvrability given their smaller head compared to body size and can chase more active prey compared to the giant blue (*B. musculus*) and fin whales (*B. physalus*) (Goldbogen et al., 2010; Nakamura et al., 2012). While employing a unique lateral suction feeding strategy, the grey whale (*Eschrichtius robustus*) has a similar scaling of the buccal cavity as rorqual whales. This species uses its baleen as a comb to extract benthic invertebrates from the bottom sediments (Berta et al., 2016). It has a mostly flat rostrum, with shorter baleen (Bouetel, 2005). Finally, the monotypic pygmy right whale (*Caperea marginata*) follows a unique allometric pattern, as it more closely approximates the trajectory of skim feeders or lunge feeders depending on the metric used. Lack of observational data for this species makes it impossible to formulate a definitive hypothesis based on its feeding mode (Werth et al., 2018).

While the effects of allometry on the feeding adaptations of baleen whales have been explored at length, very little is known on the influence of heterochronic process on their evolution and diversification. The only study to consider this topic in depth is Tsai and Fordyce (2014a). The authors compared the skulls of three prenatal and three adult specimens of pygmy right whale to prenatal and adult specimens of rorquals (sei whale and humpback whale – *Megaptera novaeangliae*) using pictures and drawings. By qualitatively comparing the specimens and conducting 2D geometric morphometric analyses, they determined that the pygmy right whale presents a paedomorphic skull shape, as the adults are similar in shape to the prenatal specimens of the species, while rorquals express peramorphic traits, given their highly different adult skull morphologies compared to their younger specimens, drawing a connection between these diverging developmental trends and the different feeding modes used by these taxa. However, this study is limited by sample size and methodology employed. Because the comparisons lacked closely related living or fossil taxa and did not use phylogenetic comparative methods to estimate ancestral states, they were unable to assess the heterochronic process in the appropriate evolutionary context (McNamara, 1986; McKinney, 1988). Therefore, while the small size of the pygmy right whale compared to all other modern baleen whales (Davies and Guiler, 1957; Pavey, 1992) could suggest that this taxon indeed has paedomorphic traits, the peramorphism of rorquals cannot be assessed by comparing them to the size and shape of *Caperea* (Gould, 1977; McNamara, 1986).

Given the hypothesised connection to the evolution of different feeding adaptions in modern Mysticeti, a quantitative analysis of skull development and allometry is needed to determine the underlying developmental changes that enabled these major dietary transitions, as well as the rapid increase in head and body size that occurred throughout this lineage (Tsai and Kohno, 2016; Fordyce and Marx, 2018). In this paper, we use 3D geometric morphometrics (GM) and phylogenetic comparative methods to identify the heterochronic processes involved in the evolution of baleen whale skull shape. We hypothesize that cranial morphology in the early fetal stages will be similar across living baleen whales, representing the phylotypic stage of the funnel developmental model (Piasecka et al., 2013; Tsai and Fordyce, 2014b). This limited diversity of shape in early development would reflect their common ancestry and the constraints posed by baleen and other skull traits related to filter feeding. We expect that disparity will increase postnatally as taxa develop unique skull shapes related to their diverse different feeding modes. We will also directly test the hypothesis by Tsai and Fordyce (2014a) that *Caperea* and rorqual whales present paedomorphic and peramorphic growth respectively compared to other mysticetes.

## 2 Materials and Methods

### 2.1 Dataset composition

The dataset is composed of 61 specimens (46 Mysticeti and 15 Odontoceti) housed in international museum collections at different ontogenetic stages, from embryos (<2 months post conception) to adults. The baleen whale sample covers the entire diversity of the clade, with at least two specimens representing five of the six living genera: *Balaena* (bowhead whales, Balaenidae), *Balaenoptera* (rorqual whales such as minke whales and the blue whale, Balaenopteridae), *Caperea* (pygmy right whale, Neobalaenidae), *Eschrichtius* (grey whale, Balaenopteridae), *Megaptera* (humpback whale, Balaenopteridae). The toothed whales were used as an outgroup in the phylogenetic comparative methods. The outgroup is composed of by five specimens of three genera representing the diversity of odontocetes (*Kogia* – pygmy sperm whales, Physeteridae; *Phocoena* – porpoises, Phocoenidae; *Lagenorhynchus* – spinner and spotted dolphins, Delphinidae).

All analyses were conducted at genus level due to the overall morphological and ecological similarity of mysticetes species, particularly in the prenatal stages (Bannister, 2018; Lanzetti, 2019), which makes it difficult to discriminate species in museum collections where outdated taxonomic names are commonly used and where associated molecular data is rare (McGowen et al., 2020). As the genus *Balaenoptera* is highly diverse and much more specious compared to the other living mysticetes (Bannister, 2018), it is represented in our sample by more specimens. We thus divided this genus in two easily identifiable taxonomic units according to recent phylogenetic analysis (McGowen et al., 2020) as well as body size: the smaller minke whales (*B. acutorostrata* and *B. bonaerensis*) and the larger sei, blue and fin whales (*B. borealis, B. musculus* and *B. physalus*).

Specimens were binned into four growth stages: early fetus (from embryonic stages to 50% of gestation time), late fetus and neonate (from 50% gestation time to birth, including the first year of life), juvenile (from second year of life to sexual maturity), and adult (after sexual maturity). This was done to ensure that minor errors in the assessment of the age of the specimens would not undermine the results, as, for most species, basic information such as the length at birth is unknown or contrasting information have been published in the literature (Frazer and Huggett, 1973; Berta et al., 2016; Lanzetti et al., 2020). Approximate age of the specimens was determined based on total length. This was either extracted from the literature or collection metadata or measured directly for specimens in fluid collections. If no information was available, body length was estimated using bizygomatic width of the skull, which is considered a reliable method to reconstruct body size in Cetacea (Pyenson and Sponberg, 2011). For prenatal specimens, the approximate age as a proportion of gestation time was reconstructed using information from the literature for each taxon and growth curves (Ohsumi et al., 1958; Laws, 1959; Tomilin, 1967; Frazer and Huggett, 1973; Masaki, 1979; Rice, 1983; Reese et al., 2001; Lanzetti, 2019) following the methodology used in Lanzetti et al. (2020). For postnatal specimens, they were defined as juveniles if their length was less than the estimated size at sexual maturity for that species according to published data (e.g. Tomilin, 1967; Rice, 1983; Bannister, 2018). Details on the specimens used in the study, including taxonomic and age assignment, can be found in Supplementary Table S1.

### 2.2 Shape data acquisition and rendering

Skulls for disarticulated osteological specimens and fluid preserved samples were digitized using medical-grade and high-resolution X-ray Computed Tomography (CT) and diffusible iodine-based contrast-enhanced X-ray CT (diceCT) (Lanzetti and Ekdale, 2021) in local institutions. Whole osteological specimens were digitized using a hand-held surface scanner (Creaform GoSCAN!) or photogrammetry, depending on availability. Digitization mode is listed for each specimen in Supplementary Table S1. Parameters and instruments used for CT scanning are listed in Supplementary Table S2, along with the resolution of each scan and source of the data if it was not scanned primarily for this project. Light surface scans were performed at 0.2-0.5 mm resolution depending on the size of the specimen.

All CT data were first imported in ImageJ (Schneider et al., 2012) to be cropped and adjusted for brightness/contrast where needed. Larger high-resolution stacks were scaled down by a factor of 2 in all 3 dimensions (i.e., binning 2×2×2), reducing the image size and limiting the number of slices to a maximum of 1500 to aid with segmentation. Images were then exported as 16-bit TIFF images stacks for segmentation in Avizo 2020.3 (Thermo Fisher Scientific Inc.). Skulls of disarticulated osteological specimens were digitally reconstructed by segmenting bone element. The manually segmented skulls were exported as OBJ mesh files for further processing. For light surface scans, the dorsal and ventral surface were digitized separately were merged in VXElements 8.1 (Creaform Inc.) and then exported as a single PLY mesh file. For photogrammetry, ReCap Photo (Autodesk Inc.) was used to create the 3D model from the images.

All surface files were then imported in Geomagic Wrap 2017 (3D Systems Inc.) for processing and cleaning. Disarticulated osteological specimens were reconstructed by aligning them to a model of a skull of similar length and species. For all meshes, holes were filled, spikes were removed, and a quick smooth was performed. All resulting surfaces were sampled to 1,500,000 triangles to ensure a consistent visualization for landmarking and then exported as PLY files.

### 2.3 Landmarking and data preparation

To quantify skull shape, 64 Type I and Type II single points landmarks (Bookstein, 1991) and 43 semilandmark curves were placed on the surfaces using Stratovan Checkpoint (Supplementary Figure S1, Table S3), and the coordinates exported in PTS format to be analysed.

The landmark configuration was chosen to capture all details of cranial shape while being identifiable in all taxa and at all growth stages represented in the dataset. Landmarks and curves that were unidentifiable due to specimen damage or because that bone or feature is absent at a specific developmental stage, were marked as ‘missing’ during landmarking in Checkpoint, which automatically assigns them a coordinate of ‘-9999’ on all three axes.

Landmarks and curves were imported into R (R Core Team, 2021) for analyses. Of the 45 Mysticeti specimens, the focus of this study, 35 out of 46 had at least one fixed landmark or curved identified as ‘missing’. Of these, 21 had “absent” structures, which in 19 specimens was the interparietal, as this bone is not visible in most baleen whale species after birth (Nakamura et al., 2016). Therefore, only 14 specimens of the samples had genuinely damaged features that were estimated before performing further analyses. The ‘fixLMtps’ function from the R package ‘Morpho’ v. 2.9 (Schlager, 2013; Schlager, 2017) was used to estimate missing landmarks by mapping weighted averages onto the missing specimen from three similar, complete configurations. Estimated landmarks are then added to each deficient configuration (Schlager, 2017). The deformation is performed by a thin-plate-spline interpolation calculated using the available landmarks (Bookstein, 1991; Schlager, 2017). For bones absent due to developmental age rather than specimen damage, all landmarks and semilandmarks associated with that structure were placed in a single “zero-area” point adjacent to its position in other taxa, following Bardua et al. (2019). List of specimens with fixed LMs or curves marked as absent is available on GitHub at https://github.com/AgneseLan/baleen-allometry.

### 2.4 Data analysis

The curves were resampled to set the same number of semilandmark points for each in all specimens using the package ‘SURGE’ (Felice, 2021), and slid on the surface to minimize bending energy (Bardua et al., 2019). After resampling the curve semilandmark, the final dataset used for analysis is composed of 462 points, of which 64 are fixed landmarks and 398 are semilandmarks. Procrustes superimposition (GPA) was performed using the ‘gpagen’ function in ‘geomorph’ v. 4.0.1 (Baken et al., 2021). This allows all specimens to be aligned and scaled, and to extract centroid size (CS) for each configuration, which will be used a measure of skull size in analyses of allometry. Age and taxonomic information were imported separately to be used as covariates. Graphics of plots was improved using the ‘ggplot2’ package v. 3.3.5 (Wickham, 2016). Code used and necessary data to repeat the analyses is available in a dedicated GitHub repository (https://github.com/AgneseLan/baleen-allometry).

#### 2.4.1 Shape variation and general allometry analysis

Variation in skull shape across the entire dataset was assessed by performing a PCA with the ‘gm.prcomp’ function as implemented in ‘geomorph’. Linear regressions for each of the first two components versus major covariates (e.g., size, taxon, growth category) were performed using the ‘lm’ function in base R to help interpret the distribution of data in the PCA plot. As size, summarized as log(CS), was found to be significantly correlated with scores for both PC1 and PC2, we used ‘procD.lm’ in ‘geomorph’ to reconstruct the common allometric regression, and then performed a PCA again on the shape residuals, in order to assess skull shape variation independent of allometric effects. Since this study is focussed on Mysticeti, we also plotted the PCAs including only baleen whale specimens, and calculated extreme shapes for this group only using the ‘shape.predictor’ function in ‘geomorph’.

#### 2.4.2 Morphological disparity and clustering

A morphological disparity analysis using the ‘morphol.disparity’ function in ‘geomorph’ was performed to assess differences between genera, both on raw shapes and common allometry residuals. If rorqual whales undergo more significant shape changes during growth than other taxa, we would expect them to have higher disparity also when the effect of size is considered, especially given the more complete growth sequence available for this group. To the same end, we performed a clustering analysis to quantitatively test if skulls of similar age of different species are more similar to each other than to the adult of the corresponding taxon. We used the ‘hclust’ function with Ward’s method (Goswami and Polly, 2010) from the ‘stats’ package to plot a dendrogram of the clusters based on Procrustes Distances (PD) among specimens. We further applied k-means clustering directly on the shape coordinate data as implemented in the ‘LloydShapes’ function from the ‘Anthropometry’ package (Vinué, 2017). The clustering analysis was performed twice, once on the entire dataset and once on a separate GPA alignment of Mysticeti. The value of k was set to 10 for the entire dataset and five for the Mysticeti-only analysis based on the results of the initial PD clustering.

#### 2.4.3 Allometry analysis by genus and ancestral state reconstruction

As shown in previous studies of Cetacea (e.g. Groves et al., 2021; Lanzetti et al., 2022), skull allometry can vary significantly across taxa. Therefore, we first tested for differences in allometry among genera and obtained taxon-specific regression parameters. We again used the ‘procD.lm’ functions, adding genus as a covariate, to reconstruct the separate allometric regressions, and tested pairwise differences between the slopes using ‘pairwise’ in the package ‘RRPP’ v. 1.1.2 (Collyer and Adams, 2021). We then used these extracted regression parameters (slopes and intercepts) to estimate the ancestral allometries and reconstruct the polarities of any heterochronic shifts (Alberch et al., 1979) following Morris et al. (2019) and Lanzetti et al. (2022). Using the phylogeny from McGowen et al. (2020), we calculated ancestral slope and intercepts parameters for the nodes with the ‘fastAnc’ function and mapped the character changes on the tree with the ‘contMap’ function, both implemented in ‘phytools’ v. 1.0 (Revell, 2012). The package ‘emmeans’ v1.7.2 (Lenth, 2022) was used to asses significant changes in the allometric slopes between ancestral nodes and extant taxa.

Finally, in order to better visualize the changes in ontogeny in the ancestral nodes within Mysticeti, we generated ancestral shapes for the prenatal and postnatal stages separately by conducting a phylogeny corrected PCA (phyloPCA) in ‘geomorph’. We first calculated the mean shape for each taxon for the prenatal (early fetal and late fetal/neonate) and postnatal (juvenile and adult) stages and then used them to conduct two separate phyloPCAs. The ancestral shapes calculated for each of the internal nodes used only for qualitative comparisons, as they are based on multiple assumptions (specimens at different growth stages, different species, etc.).

## 3 Results

### 3.1 Skull shape variation in ontogeny and phylogeny

The PCA plot for the entire dataset demonstrates a clear distinction between the two major extant clades of Cetacea (Supplementary Figure S2). PC1 (45.41% of variation) is highly correlated with phylogeny (Supplementary Table S4), with Odontoceti occupying the positive end of the axis and Mysticeti the negative end. PC2 (23.85%) instead describes changes in ontogeny, with early fetal specimens characterized by short rostra and low levels of telescoping – the overlapping of neurocranial bones and relative posterior movement of nasals typical of Cetacea (Roston and Roth, 2019) – on the positive side and adult specimens with longer rostra and more prominent telescoping on the negative one. Both axes are also highly correlated with specimen size, hence we proceeded in conducting a PCA analysis on the residuals of the common allometric model (see section 3.4 for additional details on allometric analyses). In the allometry-corrected PCA, there is again strong separation of Odontoceti and Mysticeti on PC1 (50.8%), but differences in growth stage are no longer evident on PC2 (11.9%) (Supplementary Figure S3, Table S4).

Zooming in on Mysticeti, there is marked separation of families with different feeding modes whether size is taken into account or not (Supplementary Table S5). In the raw PCA (Figure 1A), rorqual whales and the closely related *Eschrichtius* occupy the more positive side of PC1, while *Caperea* plots on the extreme negative side in line with the skim feeder *Balaena*. A strong ontogenetic gradient is visible diagonally, with adult occupying the most negative ends of both axes. In the residuals dataset (Figure 1B), differences among taxa are visible on both PC1 and PC2, with rorqual whales occupying the central portion of the morphospace and the other taxa scattered around them, with a stronger difference present between *Caperea* and *Balaena*. In both plots, there are consistent differences among taxa at all growth stages, reflecting skull shape variation associated with different feeding modes and suggesting that morphological differences among taxa appear early in ontogeny.

**Figure 1.**
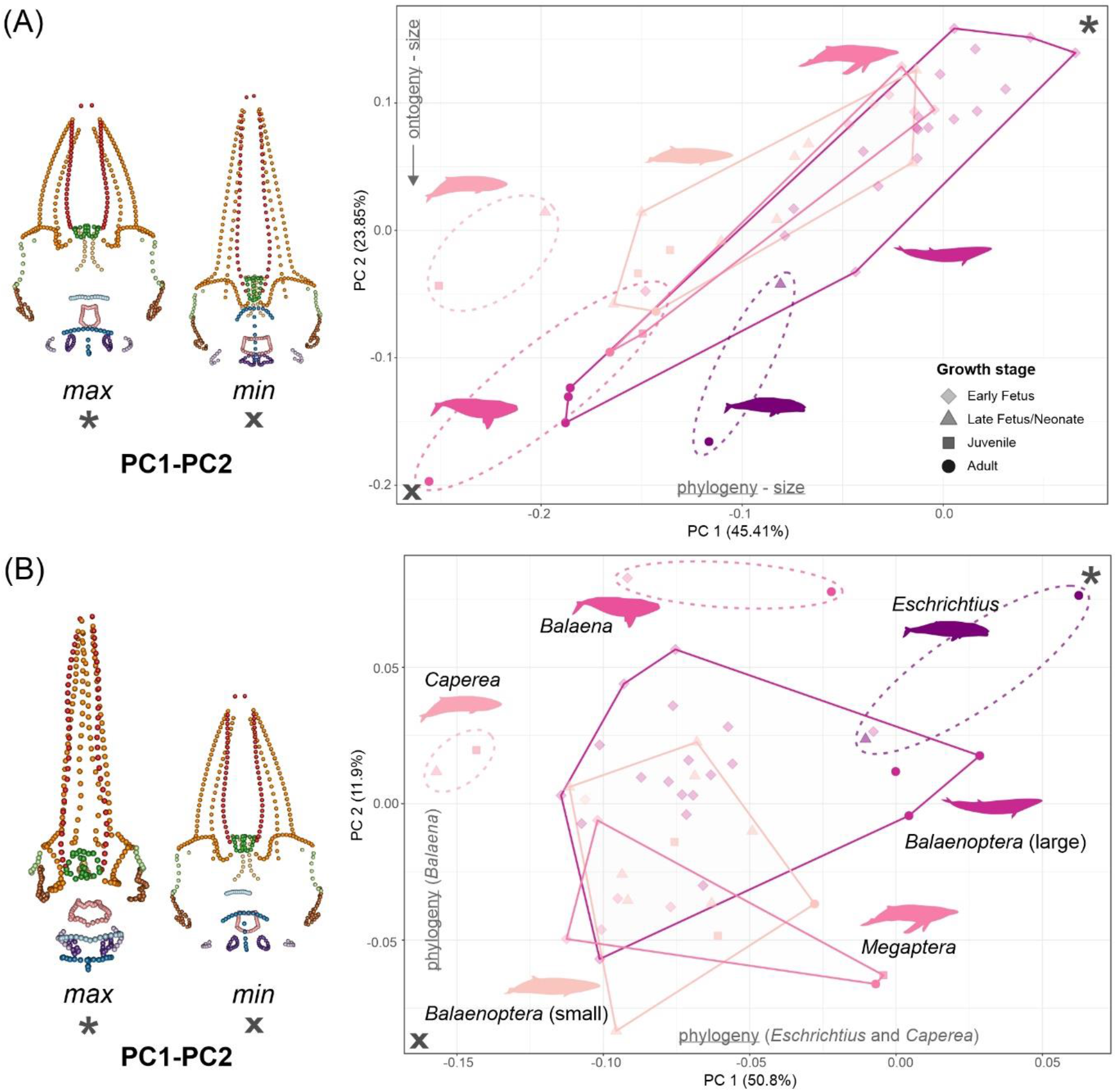
Skull ontogenetic morphospace (PCA), showing the shape variation through ontogeny and phylogeny in Mysticeti, with a significant separation of the skim feeding taxa (*Balaena* and *Caperea*) from Balaenopteridae. (A) Raw data; (B) Size-corrected using common allometry model residuals. The landmark plots on the left represent the morphological extremes at the maximum (asterisk) and minimum (cross) values of PC1 and PC2 for each plot, in dorsal view. Fixed landmarks and semilandmarks are represented in different colours to highlight shape changes in different regions of the skull (premaxilla: red, maxilla: dark orange, palatine: light orange, nasals: dark green, orbital process: light green, interparietal: light blue, supraoccipital: dark blue, squamosal: brown, exoccipital: light purple, occipital condyles: dark purple, basioccipital: pink). The factors that significantly explain shape distribution are reported on each axis. See Supplementary Tables S4-S5 for details, PCAs for full dataset in Supplementary Figure S2-S3.

### 3.2 Disparity in shape and size

In the morphological disparity analysis conducted on both raw data and allometric residuals, we found significantly different levels of disparity between mysticetes and odontocetes, as well as among the toothed whales when the effects of size are taken into account (Supplementary Table S6-S7). Among baleen whales, however, while there is a significant difference in disparity between rorqual taxa (humpback whale, minke whales, larger rorquals) and *Balaena* and *Caperea* in the raw dataset, this is not recovered when the allometric residuals are analysed, indicating that baleen whale disparity is driven by size-related shape variation (Table 1). The significant difference found in the raw dataset are likely correlated with a larger size range available for rorqual taxa, but overall, the amount of skull shape variation in ontogeny is comparable across all taxa, indicating that rorquals do not undergo a relatively higher degree of skull shape change during ontogeny.

**Table 1.** Disparity in skull shape among Mysticeti genera, in raw data and allometry residuals. Difference between Procrustes distances is reported for each pairwise comparison. Significant differences (p<0.05) are in bold.

| <u>Raw data</u> | *Balaena* | *Balaenoptera*<br>*a* (large) | *Balaenoptera*<br>*a* (small) | *Caperea* | *Eschrichtius* | *Megaptera* |
| --- | --- | --- | --- | --- | --- | --- |
| *Balaena* |  | **0.064** | **0.066** | 0.006 | 0.053 | **0.064** |
| *Balaenoptera*<br>(large) | **0.064** |  | 0.001 | **0.059** | 0.011 | 0.001 |
| *Balaenoptera*<br>(small) | **0.066** | 0.001 |  | **0.060** | 0.013 | 0.002 |
| *Caperea* | 0.006 | **0.059** | **0.060** |  | 0.047 | 0.058 |
| *Eschrichtius* | 0.053 | 0.011 | 0.013 | 0.047 |  | 0.011 |
| *Megaptera* | **0.064** | 0.001 | 0.002 | 0.058 | 0.011 |  |
| <u>Allometry</u><br><u>residuals</u> | *Balaena* | *Balaenoptera*<br>*a* (large) | *Balaenoptera*<br>*a* (small) | *Caperea* | *Eschrichtius* | *Megaptera* |
| *Balaena* |  | 0.023 | 0.023 | 0.019 | 0.022 | 0.022 |
| *Balaenoptera*<br>(large) | 0.023 |  | 0.0003 | 0.042 | 0.001 | 0.001 |
| *Balaenoptera*<br>(small) | 0.023 | 0.0003 |  | 0.042 | 0.001 | 0.001 |
| *Caperea* | 0.019 | 0.042 | 0.042 |  | 0.041 | 0.041 |
| *Eschrichtius* | 0.022 | 0.001 | 0.001 | 0.041 |  | 0.0004 |
| *Megaptera* | 0.022 | 0.001 | 0.001 | 0.041 | 0.0004 |  |

### 3.3 Specimens clustering highlights ontogenetic differences

The clustering analysis confirmed the patterns observed both in the PCA and in the disparity analyses, with morphological differences connected to different feeding modes appearing early in ontogeny, and consistent changes in shape during development among taxa. The baleen and toothed whales form distinct clusters, but Odontoceti specimens lack a clear pattern in their distribution (Supplementary Figure S4). Among Mysticeti, two main clusters are formed, one with early fetal and late fetal specimens and the second with neonates, juvenile and adults (Figure 2). In the prenatal cluster, specimens generally form small taxon-specific clusters. In the postnatal cluster, phylogeny clearly has a stronger influence, with neonates and adults of each species always plotting close to each other. Neonates and adults of rorquals plot close together in this cluster, as do the pygmy right whale specimens, as expected for a neonate and an adult. One major exception to this pattern is the grey whale (*Eschrichtius*), with the neonate plotting in the prenatal cluster away from the adult. This lack of clustering suggests that the grey whale may have an unusual ontogeny connected to its unique benthic suction feeding mode, though additional specimens need to be included to confirm this observation. The fetal specimen of *Balaena* plots in the postnatal cluster along with the adult, likely due to the marked skull shape difference between this taxon and other included in the dataset. Results of the clustering analysis are also supported by the k-means analysis pattern (Figure 2, Supplementary Figure S4).

**Figure 2.**
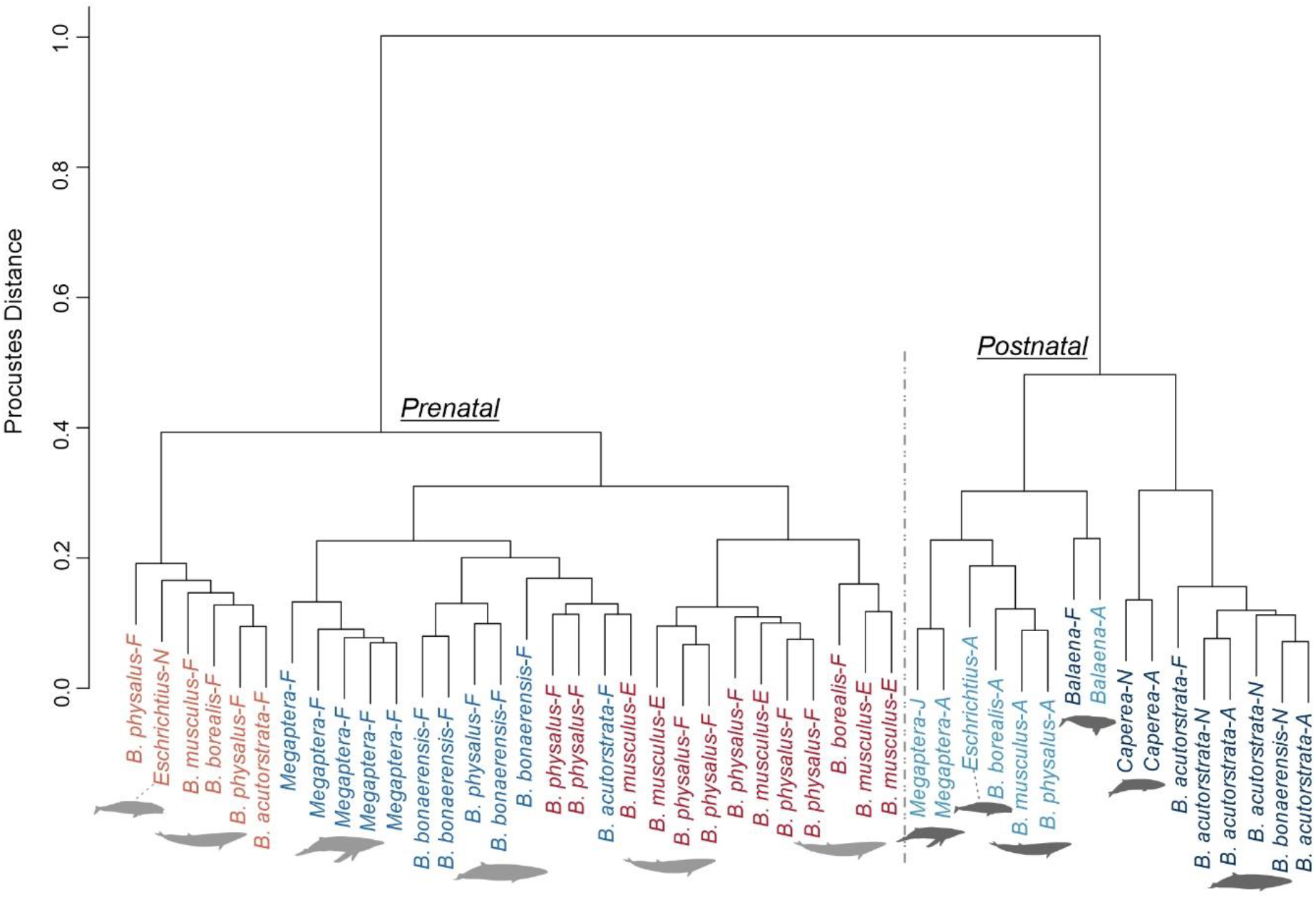
Clustering analyses on Mysticeti, represented as dendrogram of Procrustes distances. Tip labels coloured according to the results of the k-mean clustering on shape data (k=5). A prenatal and postnatal cluster can be identified and are labelled on the dendrogram. Specimens are labelled with the species names and their ontogenetic age (E = embryo, F = fetus, N = neonate, J = juvenile, A = adult). The silhouettes indicate the position of major clusters of each taxon (light grey = prenatal cluster, dark grey = postnatal cluster).

### 3.4 Conserved allometric patterns in baleen whales

Based on the distribution of shape variation in the PCA plots, as well as the consistent clustering by growth stages, we expected to find distinct allometric patterns for each taxon, but only significant differences in growth between lineages with distinct feeding modes and skull morphologies. Large and small *Balaenoptera* species and *Megaptera* display similar growth trends, while the other taxa with more divergent morphologies (*Eschrichtius, Balaena, Caperea*) are characterized by a slightly higher slope (Supplementary Figure S5, Table S8). There were no significant pairwise absolute or angular difference between slopes in the mysticete dataset, potentially reflecting the limited sample sizes available for most taxa. In terms of trajectory length, only that of the large *Balaenoptera* species is significantly longer than minke and humpback whales, likely due to their larger adult size (Supplementary Table S8).

In order to assess which heterochronic processes underlie mysticete cranial evolution of the clade, we estimated the ancestral slope and intercept values for all the taxa in the dataset. We found an inverse relationship between the slopes and intercepts values, with increases in slope accompanied by a reduction in intercept value and vice versa (Supplementary Figure S6A-B). The toothed whales used as outgroup appear to have a significantly lower growth rate than Mysticeti, but the difference is small overall and may reflect the smaller size of the odontocete taxa (Supplementary Figure S7, Table S8-S9). Looking specifically among baleen whales, we found a consistent pattern in line with our hypothesis (Figure 3). Skim feeders *Balaena* and *Caperea* have slightly accelerated growth compared to their ancestor (nodes 1 and 2), with differences between the nodes and *Balaena* also being significant. Rorquals overall have decelerated growth, in contrast with what was hypothesized in Tsai and Fordyce (2014a). The minke whales (small *Balaenoptera*) and *Megaptera* have a significantly slower growth compared to the ancestor of the group (node 3). The grey whale is the only taxon to show significantly accelerated growth among Balaenopteridae.

**Figure 3.**
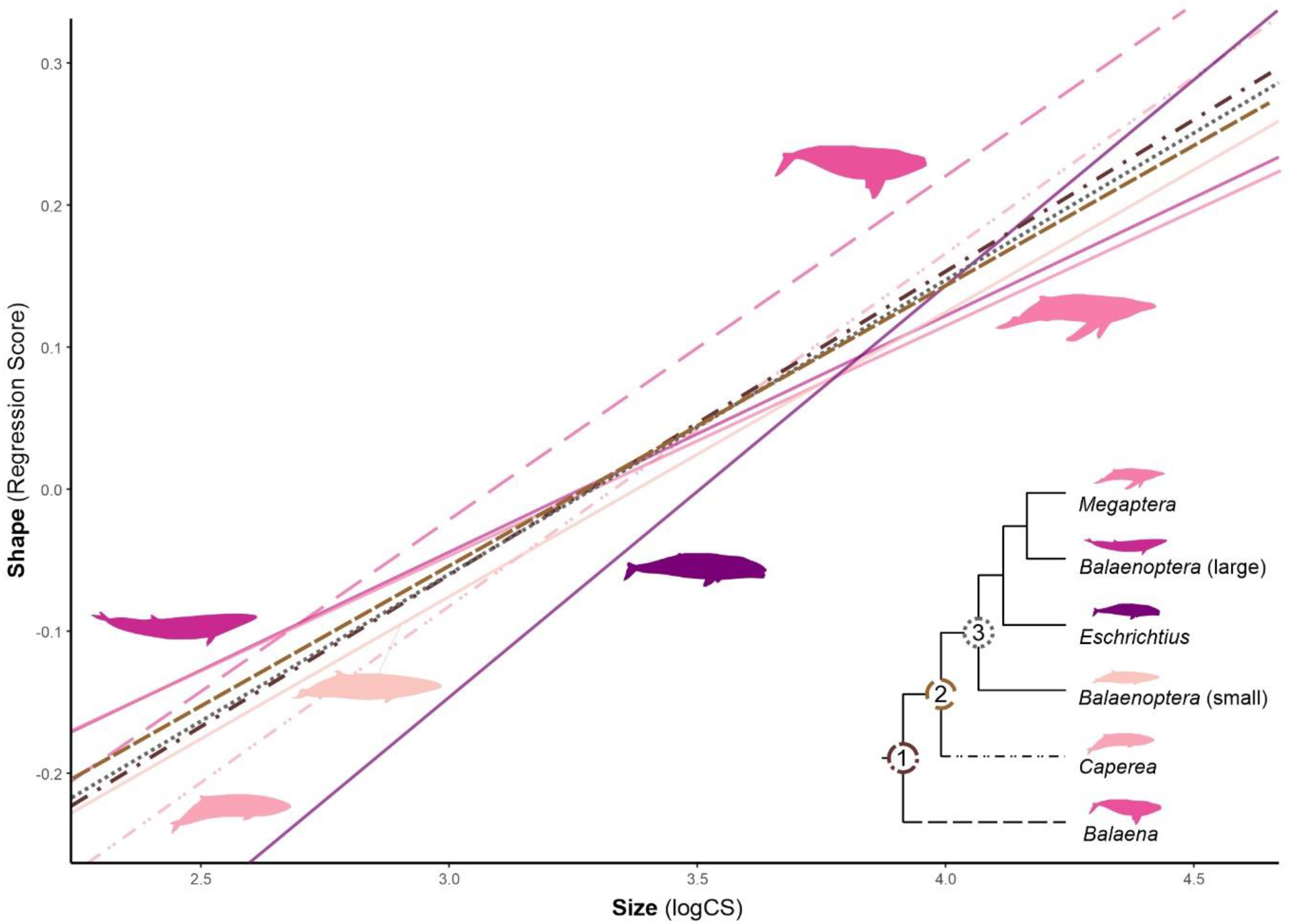
Allometric trajectories of skull shape ontogeny in Mysticeti and their ancestors. Balaenopteridae with the exception the grey whale *Eschrichtius* (solid lines) have lower slopes relative to their ancestor (node 3), indicating a paedomorphic shift. Both skim feeding taxa (*Balaena* and *Caperea*, long dash and dot-dot-dash lines) instead present a peramorphic trend relative to the ancestors (nodes 1 and 2). For full results on extant taxa see Supplementary Figure S5-Table S8, on ancestral state reconstruction see Supplementary Figure S6-S7-Table S9.

## 4 Discussion

The unique cranial shape of mysticetes facilitates their proficiency in pelagic filter feeding (Berta et al., 2016), allowing them to exploit a niche that in the past has been occupied by a variety of lineages, from armoured fishes to marine reptiles The unique cranial shape of mysticetes facilitates their proficiency in pelagic filter feeding (Berta et al., 2016), allowing them to exploit a niche that in the past has been occupied by a variety of lineages, from armoured fishes to marine reptiles (Kelley and Motani, 2015; Motani et al., 2015; Stiefel, 2020), and is presently shared also by different lineages of bony and cartilaginous fishes (Cavin, 2010; Hopman and Gilbert, 2014). The cranial adaptations required to house the long baleen plates of skim feeders or sustain the high water drag during lunge feeding clearly has influenced the evolution of the entire adult skull morphology (Bouetel, 2005), but has also appears to have constrained their ontogeny. Though, distinct developmental patterns characterize taxa with different feeding modes, underlining the importance of heterochronic changes in the diversification of this group.

### 4.1 Filter feeding constraints cranial shape development in baleen whales

#### 4.1.1 Distinct cranial morphology from the early fetal stages

All taxa undergo similar changes through ontogeny, including a progressive tapering and arching of the rostrum and an increase in the level of telescoping. However, specimens occupy defined areas of the morphospace according to their filter feeding strategy, independently of their ontogenetic age (Figure 1A-B). Similarly, the amount of shape variation at different developmental stages is comparable across taxa (Table 1). This suggest that the need to develop a skull shape capable of filter feeding, including replacing teeth with baleen (Lanzetti, 2019; Lanzetti et al., 2020), constrains overall cranial ontogeny and therefore any alteration in adult skull shape starts at the earliest stages of development but does not impact the overall degree of morphological change. Based on our dataset, we can reject our initial hypothesis that Mysticeti present a funnel-like ontogeny, with a phylotypic stage in the early fetal stages. It is possible, however, that embryonic stages are more similar across species, and this should be investigated further by examining other aspects of their ontogeny, such as ossification sequence.

This finding also calls into question the notion that mistakenly incorporating juvenile taxa in phylogenetic analysis might lead to misleading results, as suggested by Tsai and Fordyce (2014b), an issue particularly common in palaeontological studies. Especially if ratios instead of raw measurements are used, effectively correcting for size, it is possible to assume that at least any postnatal specimen would be assigned in the correct genus, and in *Balaenoptera* it would be possible to distinguish minke whales from large rorquals despite their overall morphological similarities. Other families with different feeding modes such as *Balaena* appear to be easily distinguished from rorquals even early in their ontogeny, easing any concern that including immature representatives of extinct taxa with different feeding adaptations would greatly influence the results of the phylogenetic analysis.

#### 4.1.2 Lack of evidence for paedomorphism in *Caperea*

Tsai and Fordyce (2014a) hypothesized that *Caperea* presents a paedomorphic skull shape based on the similarities between late fetal and neonate skulls to adults. We directly tested this hypothesis in our disparity (Table 1) and clustering analyses (Figure 2) and found no support for this hypothesis. Taxa clearly cluster in prenatal and postnatal groups, with consistent species-specific clusters being formed postnatally. The distance between the neonate and the adult of *Caperea* is similar to what is observed for species of *Balaenoptera* and between the fetus and adult of *Balaena*, as is the shape disparity between the two *Caperea* specimens and the other taxa. Although not sampled here, we expect that fetal specimens of *Caperea* and younger specimens of *Balaena* would plot in the prenatal cluster along with the Balaenopteridae.

Tsai and Fordyce (2014a) based their hypothesis on the observation that Balaenopteridae appeared to undergo a larger degree of shape change during ontogeny. We find consistent distances and shape disparity across rorqual whale species, with the only exception being the grey whale. This taxon seems to undergo a greater degree of shape change postnatally, as highlighted by its distinctive placement in the clustering analyses, with its neonate plotting in the prenatal cluster. This odd placement might be due to the lack of prenatal specimens for this taxon in the dataset, and it should be investigated further to confirm this preliminary hypothesis.

Overall, given the consistent shape development pattern highlighted in all our analyses, we conclude that all Mysticeti likely undergo similar levels of shape changes during their ontogeny, with some timing difference possibly related to their characteristics feeding modes and their influence on skull morphology (Werth et al., 2018). This reaffirms our previous hypothesis of a shared skull shape development pattern in modern baleen whales, with morphological differences already arising early in their ontogeny. Therefore, we need to look into other aspects of development to understand the origin of the disparate morphologies, feeding modes and body size present in the group today and recorded in the fossil record.

### 4.2 Heterochronic changes in allometry supported the evolution of diverse feeding modes

#### 4.2.1 Paedomorphic trend in rorqual whales

While skull shape ontogeny appears to vary little variation mysticetes, allometry varies significantly among taxa. Balaenopteridae in particular stands out for having a slower growth compared to their reconstructed ancestors (Figure 3). While this is to be expected for larger rorquals and *Megaptera* given the need to reach a larger skull size while maintaining a constant amount of shape change as we established earlier, the smaller minke whales also follow this same trend, suggesting that is the feeding mode they all share rather than body size alone driving this change in their allometry. These data allows us to reject the hypothesis of Tsai and Fordyce (2014a) that rorqual whales present peramorphic growth. Instead, all taxa in that group have significantly decelerated or paedomorphic growth compared to their ancestral estimate (Figure 4A). The larger taxa compensate for this slower growth by lengthening their developmental time, allowing them to reach their impressive body and size while managing current allometric scaling for lunge filter feeding (Goldbogen et al., 2012; Goldbogen and Madsen, 2018; Kahane-Rapport and Goldbogen, 2018; Goldbogen et al., 2019). The sei whale, blue whale and fin whale have a gestation of about 12 months, humpback whales of 11 months, and while the smaller minke whales only gestate for about around 10 months (Lanzetti et al., 2020). Additionally, larger whales also need to reach a greater body size to be considered adults, adding to their overall developmental time (Bannister, 2018).

**Figure 4.**
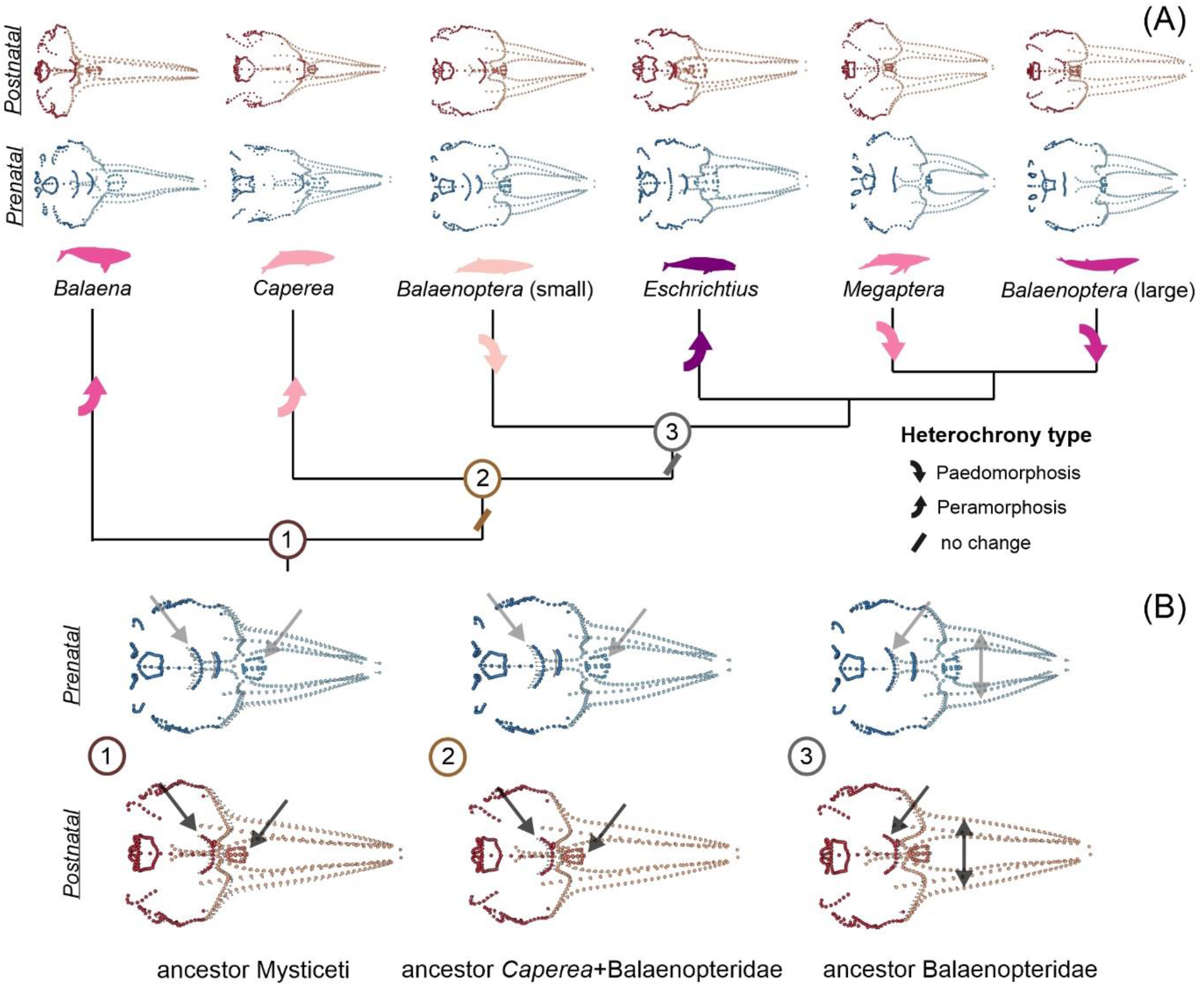
Heterochrony in Mysticeti phylogeny and related skull shape changes. (A) Phylogeny with inferred direction of heterochronic change for each node and mean shapes for prenatal (blue) and postnatal (red) stages of each taxon; (B) reconstructed prenatal and postnatal skull shapes at the ancestral nodes. Landmarks in the neurocranial region are highlighted with a darker shade, nasals have a medium shade, and the rostrum has a lighter shade. The arrows on the ancestral skull shapes point to the region where the developmental differences with modern taxa are more marked, as described in the text. Skull shapes in dorsal view (for medial view see Supplementary Figure S8), not to scale.

#### 4.2.2 Peramorphism and the evolution of skim and suction filter feeding

On the other hand, skim feeding *Balaena* and *Caperea* as well as the suction feeding *Eschrichtius* all present faster growth than the rorqual whales and are characterized by peramorphic shifts in their allometry compared to the reconstructed ancestral states (Figure 4A). While this result needs to be interpreted with caution given the limited sample size available for these taxa, it is possible that the development of certain skull features that allow them to employ skim and suction filter feeding is dependent on this peramorphic shift in development. For example, rostrum arching appears to be a feature that progressively increases during ontogeny, and therefore a taxon with a fully arched palate like *Balaena* might require a faster rate of development to reach the desired rostral shape while maintaining the correct scaling of head and body size (Werth et al., 2018). Additionally, a close relative of this taxon and also a skim feeder (*Eubalaena glacialis* – North Atlantic right whale) was documented to have a fast growth rate in the early postnatal period before maturity (Fortune et al., 2012). While this might be a strategy that evolved to ensure that claves quickly reach a large body size to avoid predation, it also helps maximize foraging efficiency by ensuring a fast development and correct scaling of the rostrum and the baleen. The suction feeding grey whale instead, while not presenting exaggerated skull features relative to other Balaenopteridae, has a unique ecology: it migrates across the Pacific Ocean from the North Pole to the breeding grounds in Mexico (Swartz, 2018). This migration is the longest recorded in baleen whales, and it has been shown to directly affect the fetal development, as it slows down while pregnant mothers make their journey from North to South, and it accelerates again when they reach the breeding grounds (Rice, 1983). This period of little to no growth might cause the overall rate to be higher in order to complete the development. While little information is available on *Caperea*, including specific on its feeding mode and its development, some of its skull traits are similar to skim feeders (Werth et al., 2018), and therefore it is possible that it shares the same peramorphic growth trend.

#### 4.2.3 Ontogeny tracks the evolution of specialized filter feeding modes in the fossil record

While allometric trends correlate well with the present feeding mode diversity in Mysticeti, the reconstructed ancestral allometries and skull shapes for both the prenatal and postnatal stages allow us to hypothesize the likely feeding modes and developmental patterns at the ancestral nodes (Figure 4B). The ancestor of all crown Mysticeti (node 1) appears to have an intermediate skull shape development between lunge and skim feeding taxa. The telescoping pattern was probably less marked in ontogeny: in the hypothetical prenatal stages the supraoccipital shield already occupied a more forward position, and the nasals were more posterior and tapered, all traits associated with later ontogenetic stages in modern taxa. The dorsal arching of the rostrum increased during development, as did rostral tapering, but not to the same degree as seen in modern skim feeding taxa (Supplementary Figure S8). This would suggest that they were capable of a less specialized form of filter feeding, usually referred to as bulk filter feeding (Berta et al., 2016). The hypothesis that at least some lineages of fossil mysticetes employed a less generalized feeding strategy is supported by the remarkable findings of fossilized stomach contents and baleen plates in Miocene specimens (Collareta et al., 2015; Marx et al., 2017). These fossils belong to the paraphyletic extinct group “Cetotheriidae†” (Berta et al., 2016). The uncertainty over the evolutionary relationships of this group is probably due to convergent evolution of similar filter feeding modes displayed in specimens found in different parts of the world, as well as to the unique mixture of traits they presented (Bisconti, 2015; Gol’din et al., 2015; Berta et al., 2016). For example, some of them had short baleen plates that closely resemble the morphology of *Caperea* (Marx et al., 2017), which has been proposed to be included as the last living representative of the group (Fordyce and Marx, 2013; Marx et al., 2019), although this is still debated (Berta et al., 2016; Gatesy et al., 2022), but lacked some morphological features seen in the modern species.

Similar fossils from the same locality have been found with fish fossilized in their stomach, hinting that they had a diet more similar to modern rorquals in contrast with the copepod-specialist pygmy right whale (Collareta et al., 2015).

The ancestor of *Caperea* and Balaenopteridae (node 2) presents similar mixed traits in rostral morphology and a comparable progression of telescoping as the ancestor of all crown Mysticeti (node 1). This suggests that the likely skim feeding specialization in *Caperea* might have evolved convergently to Balaenidae rather than being inherited by a common ancestor, accompanied by an acceleration in its development. This is in line with a recent study that analysed pattern of rostral morphology evolution across extant and extinct baleen whales (Tanaka, 2022).

Despite the presence of *Eschrichtius* in the dataset, the reconstructed ancestor of Balaenopteridae (node 3) closely matches the morphology and ontogenetic changes seen in modern rorqual whales. Particularly, between the prenatal and postnatal stages there is a stronger anterior shift of the supraoccipital shield as well as a more marked rostral tapering. While the reconstructed allometric trend is similar to the other nodes examined, the hypothesized postnatal skull shape perfectly exemplifies the concept of paedomorphism, as it appears to preserve traits such as a less anteriorly shifted supraoccipital or less arched rostrum that would be considered immature in their ancestors (node 1 and node 2). Moreover, its similarity with modern rorquals suggest that some extinct taxa might have utilized a form of lunge feeding rather than generic bulk filter feeding. While most known fossil lineages did not present traits associated with specialized feeding strategies, there is evidence that at least one *Herpetocetus morrowi*† likely employed lateral suction filter feeding to consume benthic invertebrates similar to the grey whale (El Adli et al., 2014). Given the unique allometry of *Eschrichtius*, it is possible that convergent ontogenetic shifts in these two lineages allowed for the evolution of this unique feeding strategy, and this hypothesis could be explored further in the future by reconstructing the allometry of extinct taxa with multiple specimens preserved.

In conclusion, in this novel study applying 3D morphometrics to a comprehensive comparative dataset, we were able to effectively test the role of heterochrony in different aspects of skull ontogeny in baleen whales. We were able to refute two hypotheses originally formulated using qualitative data: that Mysticeti have a funnel-like ontogeny with a conserved skull shape in early ontogeny and that *Caperea* presents paedomorphic development while Balaenopteridae have peramorphic growth. We instead conclude that Mysticeti present distinct cranial morphologies connected to their feeding modes from the early fetal stages. They then follow a shared developmental pattern, with a similar degree of shape change occurring in all taxa. While constraint in shape change appears to be the key for developing functional skull for general filter feeding, small heterochronic variation in the rate of growth and scaling of the skull can give rise to a variety of morphologies best adapted to different filter feeding modes. Changes in allometry play a key role in establishing the differences observed in the adults of each species and in maintaining the correct scaling of the head relative to body size to best develop each feeding mode. Lunge feeding Balaenopteridae underwent a paedomorphic shifts during their evolution and present a slower growth. Larger species can reach their impressive body size by having extended their developmental time. Both skim and suction filter feeders are characterized by a faster peramorphic allometric growth. These conclusions and our reconstruction of ancestral allometric trajectories and skull shapes are broadly supported by fossil evidence and by other studies on the physiology, ecology, and feeding mechanisms of modern species. Additional specimens of less represented taxa in our dataset such as the grey whale and the pygmy right whale are needed to confirm their developmental pattern and better characterize the connection between their ontogeny and their unique feeding modes. A future comparative study between Mysticeti and Odontoceti with a larger dataset will be able to clarify whether these patterns are unique to baleen whales or if they share aspects of their development with toothed whales.

