## Supplementary for "Testing heterochrony: Connecting skull shape ontogeny and evolution of feeding adaptations in baleen whales"

**Table S1. List of specimens analysed in the study *(separate ".csv" file)*.**
This table includes specimens' codes used to identify specimens in analyses, museum accession number, taxonomic information, total length and bizygomatic width, age estimate, growth category assignment, specimen preparation, scan type, as well as reference to who provided the data. Additional notes are provided in a separate sheet.

**Table S2. Details on acquisition specimens digitized using CT scanning *(separate ".csv" file)*.**
Facility, CT system model used and details on image acquisition (kV, mA, slice thickness, voxel size, number of images, output format for images) are reported, along with other details when available. Additional notes are provided in a separate sheet.

**Table S3. Landmarks (LM), curves and number of semi-landmarks (sLMs) used in this study.**
Left landmarks and curves are italicized, points on the midline are in bold. Landmarks and curves are pictured with fixed landmarks numbered in Figure S1.

| Bone | LM | Landmark definition | Curve | Curve definition | No. sLMs |
|---|---|---|---|---|---|
| maxilla | *1*, 10 | anterior end maxilla | *1-2*, 10-11 | lateral edge of rostral portion of maxilla | 25 |
| maxilla | *2*, 11 | antorbital notch - deepest point | *2-3*, 11-12 | anterior margin antorbital notch | 5 |
| maxilla | *3*, 12 | lateral edge of antorbital notch (maxilla) | *3-4*, 12-13 | lateral and posterior edge ascending process maxilla | 20 |
| maxilla | *4*, 13 | medial end ascending process of maxilla (dorsalmost point of contact with nasal/ethmoid) | — | — | — |
| maxilla | *5*, 14 | posterior end premaxilla (contact on maxilla) | *1-5*, 10-14 | lateral edge premaxilla on maxilla | 25 |
| nasal | *6*, 15 | ventral/anterior medial end nasal | *6-8*, 15-17 | ventrolateral edge of nasal | 5 |
| nasal | *7*, 16 | dorsal/posterior medial end nasal | *8-6*, 17-15 | dorsomedial edge of nasal | 5 |
| nasal | *8*, 17 | dorsal/posterior lateral end nasal | — | — | — |
| nasal | *9*, 18 | ventral/anterior lateral end nasal | — | — | — |
| premaxilla | *19*, 20 | anterior end premaxilla | — | — | — |
| frontal | *21*, 28 | ventral end preorbital process of frontal | *21-22*, 28-29 | orbital process of frontal | 7 |
| frontal | *22*, 29 | ventral end postorbital process of frontal | — | — | — |
| squamosal | *23*, 30 | anterior end supramastoid crest of squamosal | *23-24*, 30-31 | anterior edge of squamosal | 3 |
| squamosal | *24*, 31 | anterior end zygomatic process of squamosal | *24-25*, 31-32 | ventral edge of squamosal | 5 |
| squamosal | *25*, 32 | ventral end postglenoid process of squamosal | *25-26*, 32-33 | posterior edge of squamosal | 10 |
| squamosal | *26*, 33 | posterior end supramastoid crest of squamosal | *26-23*, 33-30 | dorsal edge of squamosal | 5 |
| exoccipital | *27*, 34 | lateralmost edge of exoccipital (contact -present or future- with posteriormost edge of squamosal plate) | *35-27*, 39-34 | lateral edge of exoccipital | 7 |
| exoccipital | *35*, 39 | ventral end paraoccipital process of exoccipital | — | — | — |
| occipital condyle | *36*, 40 | dorsal end occipital condyle | *37-36*, 41-40 | lateral edge of occipital condyle | 7 |
| occipital condyle | *37*, 41 | ventral end occipital condyle | *36-37*, 40-41 | medial edge of occipital condyle | 7 |
| basioccipital | *38*, 42 | posterior end of the basioccipital crest | *53-38*, 53-42 | posterior edge of basioccipital | 5 |
| maxilla | *48*, 43 | anterior end tooth row on maxilla | *48-49*, 43-44 | medial edge of tooth row on maxilla | 20 |
| maxilla | *49*, 44 | posterior end tooth row on maxilla | — | — | — |
| jugal | *50*, 45 | anterior end maxillary process of jugal (anteriormost point of contact between jugal and lacrimal/frontal) | — | — | — |
| palatine | *51*, 46 | anteriormost point of palatine crest, edge of greater palatal foramen | *51-52*, 46-47 | medial edge of palatine | 9 |
| palatine | *52*, 47 | posteriormost point of nasal spine of palatine | — | — | — |
| basioccipital | *56*, 54 | anteriormost lateral edge of basioccipital | — | — | — |
| interparietal | *57*, 59 | anterolateral edge of interparietal (contact -present or future- with frontal) | — | — | — |
| supraoccipital | *60*, 62 | lateral edge of supraoccipital (contact -present or future- with interparietal or limit of medial part of suproccipital if interparietal obscured) | — | — | — |
| basioccipital | **53** | posteriormost edge of basioccipital | *38-56*, 42-54 | lateral edge of basioccipital | 10 |

| | | | | | |
|---|---|---|---|---|---|
| basioccipital | **55** | anteriormost edge of basioccipital | *56-55*, 54-55 | anterior edge of basioccipital | 5 |
| interparietal | **58** | midpoint anterior margin of interparietal (contact with frontal) | *57-58*, 59-58 | anterior edge of interparietal (contact with frontal) | 6 |
| supraoccipital | **61** | midpoint anterior margin of supraoccipital (contact with interparietal), dorsal end of supraoccipital crest | *60-61*, 62-61 | dorsal edge of supraoccipital (contact with interparietal) | 5 |
| supraoccipital | **63** | keyhole notch, ventral end of supraoccipital crest | **61-63** | occipital crest | 6 |
| vomer | **64** | anteriormost point of contact between right and left palatal processes of maxillae or palatines (whichever most anterior) | — | — | — |

**Table S4. ANOVA analyses of the first two PC components for raw shape data and allometry residuals.** PC components obtained from the ‘gm.prcomp’ function in ‘geomorph’, ‘lm’ function in base R used to conduct ANOVA analyses. Four independent variables were tested for raw data: size (log-transformed centroid size), genera, growth category and taxonomic group (Mysticeti and Odontoceti). Size was not tested for residuals as the allometry correction eliminates the effect of size from the data. Results reported as ANOVA tables. Significant p-values (p<0.05) are in bold.

| **Raw data** | | | | | | | | | | | |
|---|---|---|---|---|---|---|---|---|---|---|---|
| PC1 | | | | | | PC2 | | | | | |
| *ANOVA table size* | | | | | | *ANOVA table size* | | | | | |
| | Df | Sum_Sq | Mean_Sq | F | Pr(>F) | | Df | Sum_Sq | Mean_Sq | F | Pr(>F) |
| *size* | 1 | 0.54812 | 0.54812 | 43.00 | **1.52E-08** | *size* | 1 | 0.30447 | 0.304474 | 47.44 | **4.19E-09** |
| Residuals | 59 | 0.75244 | 0.01275 | | | Residuals | 59 | 0.37867 | 0.006418 | | |
| *ANOVA table growth category* | | | | | | *ANOVA table growth category* | | | | | |
| | Df | Sum_Sq | Mean_Sq | F | Pr(>F) | | Df | Sum_Sq | Mean_Sq | F | Pr(>F) |
| *category* | 3 | 0.08739 | 0.029129 | 1.37 | 0.2616 | *category* | 3 | 0.41359 | 0.137865 | 29.15 | **1.48E-11** |
| Residuals | 57 | 1.21317 | 0.021284 | | | Residuals | 57 | 0.26955 | 0.004729 | | |
| *ANOVA table group* | | | | | | *ANOVA table group* | | | | | |
| | Df | Sum_Sq | Mean_Sq | F | Pr(>F) | | Df | Sum_Sq | Mean_Sq | F | Pr(>F) |
| *group* | 1 | 0.97757 | 0.97757 | 178.57 | **<2.20E-16** | *group* | 1 | 0.11859 | 0.118595 | 12.39 | **0.0008373** |
| Residuals | 59 | 0.32299 | 0.00547 | | | Residuals | 59 | 0.56455 | 0.009569 | | |
| *ANOVA table genera* | | | | | | *ANOVA table genera* | | | | | |
| | Df | Sum_Sq | Mean_Sq | F | Pr(>F) | | Df | Sum_Sq | Mean_Sq | F | Pr(>F) |
| *genera* | 8 | 1.11832 | 0.13979 | 39.89 | **<2.20E-16** | *genera* | 8 | 0.25299 | 0.031624 | 3.82 | **0.001347** |
| Residuals | 52 | 0.18224 | 0.003505 | | | Residuals | 52 | 0.43015 | 0.008272 | | |
| **Allometry residuals** | | | | | | | | | | | |
| PC1 | | | | | | PC2 | | | | | |
| *ANOVA table growth category* | | | | | | *ANOVA table growth category* | | | | | |
| | Df | Sum_Sq | Mean_Sq | F | Pr(>F) | | Df | Sum_Sq | Mean_Sq | F | Pr(>F) |
| *category* | 3 | 0.16895 | 0.056318 | 3.79 | **0.01508** | *category* | 3 | 0.023853 | 0.007951 | 2.12 | 0.1082 |
| Residuals | 57 | 0.8472 | 0.014863 | | | Residuals | 57 | 0.214197 | 0.0037578 | | |
| *ANOVA table group* | | | | | | *ANOVA table group* | | | | | |
| | Df | Sum_Sq | Mean_Sq | F | Pr(>F) | | Df | Sum_Sq | Mean_Sq | F | Pr(>F) |
| *group* | 1 | 0.85433 | 0.85433 | 311.48 | **<2.20E-16** | *group* | 1 | 0.000258 | 0.000258 | 0.06 | 0.8012 |
| Residuals | 59 | 0.16182 | 0.00274 | | | Residuals | 59 | 0.237792 | 0.00403 | | |
| *ANOVA table genera* | | | | | | *ANOVA table genera* | | | | | |
| | Df | Sum_Sq | Mean_Sq | F | Pr(>F) | | Df | Sum_Sq | Mean_Sq | F | Pr(>F) |
| *genera* | 8 | 0.88884 | 0.111105 | 45.38 | **<2.20E-16** | *genera* | 8 | 0.205556 | 0.0256945 | 41.12 | **<2.20E-16** |
| Residuals | 52 | 0.12732 | 0.002448 | | | Residuals | 52 | 0.032494 | 0.0006249 | | |

**Table S5. ANOVA analysis of taxonomic differences in Mysticeti in first two PC components for raw shape data and allometry residuals.**
PC components obtained from the ‘gm.prcomp’ function in ‘geomorph’, ‘lm’ function in base R used to conduct ANOVA analysis on generic differences among Mysticeti. Results reported as summary table to show differences from the reference taxon (*Balaena*). Significant p-values (p<0.05) are in bold.

**Raw data**

| PC1 | | | | | PC2 | | | | |
|---|---|---|---|---|---|---|---|---|---|
| Genera | Estimate | Std. Error | t value | Pr(>\|t\|) | Genera | Estimate | Std. Error | t value | Pr(>\|t\|) |
| *Balaena* (ref.) | -0.202 | 0.047 | -4.333 | **>0.001** | *Balaena* (ref.) | -0.122 | 0.059 | -2.058 | **0.046** |
| *Balaenoptera* (large) | 0.168 | 0.049 | 3.452 | **0.001** | *Balaenoptera* (large) | 0.173 | 0.062 | 2.780 | **0.008** |
| *Balaenoptera* (small) | 0.107 | 0.050 | 2.121 | **0.040** | *Balaenoptera* (small) | 0.143 | 0.064 | 2.226 | **0.032** |
| *Caperea* | -0.023 | 0.066 | -0.342 | 0.734 | *Caperea* | 0.108 | 0.084 | 1.283 | 0.207 |
| *Eschrichtius* | 0.103 | 0.066 | 1.566 | 0.125 | *Eschrichtius* | 0.018 | 0.084 | 0.218 | 0.828 |
| *Megaptera* | 0.141 | 0.053 | 2.663 | **0.011** | *Megaptera* | 0.169 | 0.067 | 2.512 | **0.016** |

**Allometry residuals**

| PC1 | | | | | PC2 | | | | |
|---|---|---|---|---|---|---|---|---|---|
| Genera | Estimate | Std. Error | t value | Pr(>\|t\|) | Genera | Estimate | Std. Error | t value | Pr(>\|t\|) |
| *Balaena* (ref.) | -0.057 | 0.026 | -2.182 | **0.035** | *Balaena* (ref.) | 0.080 | 0.018 | 4.514 | **>0.001** |
| *Balaenoptera* (large) | -0.007 | 0.027 | -0.245 | 0.807 | *Balaenoptera* (large) | -0.070 | 0.019 | -3.782 | **0.001** |
| *Balaenoptera* (small) | -0.019 | 0.028 | -0.677 | 0.502 | *Balaenoptera* (small) | -0.101 | 0.019 | -5.264 | **>0.001** |
| *Caperea* | -0.093 | 0.037 | -2.528 | **0.016** | *Caperea* | -0.065 | 0.025 | -2.568 | **0.014** |
| *Eschrichtius* | 0.083 | 0.037 | 2.247 | **0.030** | *Eschrichtius* | -0.030 | 0.025 | -1.202 | 0.236 |
| *Megaptera* | -0.014 | 0.030 | -0.483 | 0.632 | *Megaptera* | -0.124 | 0.020 | -6.132 | **>0.001** |

**Table S6. Pairwise comparison of morphological disparity in skull shape development among genera calculated with raw data.**
Disparity analysis was performed using the ‘morphol.disparity’ function in ‘geomorph’. Disparity is calculated as differences in Procrustes variances across the entire sample for each genus using the raw data dataset. P-values calculated with 1000 permutations. Significant p-values (p<0.05) are in bold.

Procrustes variances for genera

| | *Balaena* | *Balaenoptera* (large) | *Balaenoptera* (small) | *Caperea* | *Eschrichtius* | *Kogia* | *Lagenorhynchus* | *Megaptera* | *Phocoena* |
|---|---|---|---|---|---|---|---|---|---|
| | 0.091 | 0.026 | 0.025 | 0.085 | 0.038 | 0.140 | 0.069 | 0.027 | 0.068 |

Pairwise absolute differences in path distances and p-values

| | *Balaena* | *Balaenoptera* (large) | *Balaenoptera* (small) | *Caperea* | *Eschrichtius* | *Kogia* | *Lagenorhynchus* | *Megaptera* | *Phocoena* |
|---|---|---|---|---|---|---|---|---|---|
| *Balaena* | | **0.064** | **0.066** | 0.006 | 0.053 | 0.049 | 0.022 | **0.064** | 0.023 |
| p-value | | ***0.033*** | ***0.030*** | *0.852* | *0.180* | *0.129* | *0.522* | ***0.047*** | *0.479* |
| *Balaenoptera* (large) | **0.064** | | 0.001 | **0.059** | 0.011 | **0.113** | **0.043** | 0.001 | **0.042** |
| p-value | ***0.033*** | | *0.942* | ***0.047*** | *0.679* | ***0.001*** | ***0.030*** | *0.962* | ***0.020*** |
| *Balaenoptera* (small) | **0.066** | 0.001 | | **0.060** | 0.013 | **0.115** | **0.044** | 0.002 | **0.043** |
| p-value | ***0.030*** | *0.942* | | ***0.042*** | *0.685* | ***0.001*** | ***0.030*** | *0.906* | ***0.028*** |
| *Caperea* | 0.006 | **0.059** | **0.060** | | 0.047 | 0.055 | 0.016 | 0.058 | 0.017 |
| p-value | *0.852* | ***0.047*** | ***0.042*** | | *0.216* | *0.078* | *0.622* | *0.066* | *0.590* |
| *Eschrichtius* | 0.053 | 0.011 | 0.013 | 0.047 | | **0.102** | 0.031 | 0.011 | 0.031 |
| p-value | *0.180* | *0.679* | *0.685* | *0.216* | | ***0.005*** | *0.318* | *0.706* | *0.313* |
| *Kogia* | 0.049 | **0.113** | **0.115** | 0.055 | **0.102** | | **0.070** | **0.113** | **0.071** |
| p-value | *0.129* | ***0.001*** | ***0.001*** | *0.078* | ***0.005*** | | ***0.009*** | ***0.001*** | ***0.002*** |

| | | | | | | | | | |
|---|---|---|---|---|---|---|---|---|---|
| *Lagenorhynchus* | 0.022 | **0.043** | **0.044** | 0.016 | 0.031 | **0.070** | | 0.042 | 0.001 |
| p-value | *0.522* | ***0.030*** | ***0.030*** | *0.622* | *0.318* | ***0.009*** | | *0.062* | *0.969* |
| *Megaptera* | **0.064** | 0.001 | 0.002 | 0.058 | 0.011 | **0.113** | 0.042 | [redacted] | 0.041 |
| p-value | ***0.047*** | *0.962* | *0.906* | *0.066* | *0.706* | ***0.001*** | *0.062* | [redacted] | *0.058* |
| *Phocoena* | 0.023 | **0.042** | **0.043** | 0.017 | 0.031 | **0.071** | 0.001 | 0.041 | |
| p-value | *0.479* | ***0.020*** | ***0.028*** | *0.590* | *0.313* | ***0.002*** | *0.969* | *0.058* | |

**Table S7. Pairwise comparison of morphological disparity in skull shape development among genera calculated with allometry residuals.**

Disparity analysis was performed using the 'morphol.disparity' function in 'geomorph'. Disparity is calculated as differences in Procrustes variances across the entire sample for each genus using the allometry residuals. P-values calculated with 1000 permutations. Significant p-values (p<0.05) are in bold.

Procrustes variances for genera

| | *Balaena* | *Balaenoptera* (large) | *Balaenoptera* (small) | *Caperea* | *Eschrichtius* | *Kogia* | *Lagenorhynchus* | *Megaptera* | *Phocoena* |
|---|---|---|---|---|---|---|---|---|---|
| | 0.039 | 0.016 | 0.016 | 0.058 | 0.017 | 0.109 | 0.063 | 0.017 | 0.052 |

Pairwise absolute differences in path distances and p-values

| | *Balaena* | *Balaenoptera* (large) | *Balaenoptera* (small) | *Caperea* | *Eschrichtius* | *Kogia* | *Lagenorhynchus* | *Megaptera* | *Phocoena* |
|---|---|---|---|---|---|---|---|---|---|
| *Balaena* | | 0.023 | 0.023 | 0.019 | 0.022 | **0.071** | 0.024 | 0.022 | 0.013 |
| p-value | | *0.245* | *0.266* | *0.457* | *0.403* | ***0.027*** | *0.331* | *0.322* | *0.559* |
| *Balaenoptera* (large) | 0.023 | [redacted] | 0.0003 | 0.042 | 0.001 | **0.094** | **0.047** | 0.001 | **0.036** |
| p-value | *0.245* | [redacted] | *0.976* | *0.066* | *0.968* | ***0.001*** | ***0.017*** | *0.915* | ***0.023*** |
| *Balaenoptera* (small) | 0.023 | 0.0003 | | 0.042 | 0.001 | **0.093** | **0.047** | 0.001 | **0.036** |
| p-value | *0.266* | *0.976* | | *0.077* | *0.972* | ***0.001*** | ***0.021*** | *0.941* | ***0.028*** |
| *Caperea* | 0.019 | 0.042 | 0.042 | [redacted] | 0.041 | 0.051 | 0.005 | 0.041 | 0.006 |
| p-value | *0.457* | *0.066* | *0.077* | [redacted] | *0.172* | *0.059* | *0.844* | *0.096* | *0.771* |
| *Eschrichtius* | 0.022 | 0.001 | 0.001 | 0.041 | | **0.092** | 0.046 | 0.0004 | 0.035 |
| p-value | *0.403* | *0.968* | *0.972* | *0.172* | | ***0.007*** | *0.093* | *0.990* | *0.144* |
| *Kogia* | **0.071** | **0.094** | **0.093** | 0.051 | **0.092** | [redacted] | **0.047** | **0.092** | **0.057** |
| p-value | ***0.027*** | ***0.001*** | ***0.001*** | *0.059* | ***0.007*** | [redacted] | ***0.038*** | ***0.001*** | ***0.008*** |
| *Lagenorhynchus* | 0.024 | **0.047** | **0.047** | 0.005 | 0.046 | **0.047** | | **0.045** | 0.011 |
| p-value | *0.331* | ***0.017*** | ***0.021*** | *0.844* | *0.093* | ***0.038*** | | ***0.020*** | *0.591* |
| *Megaptera* | 0.022 | 0.001 | 0.001 | 0.041 | 0.0004 | **0.092** | **0.045** | [redacted] | 0.035 |
| p-value | *0.322* | *0.915* | *0.941* | *0.096* | *0.990* | ***0.001*** | ***0.020*** | [redacted] | *0.054* |
| *Phocoena* | 0.013 | **0.036** | **0.036** | 0.006 | 0.035 | **0.057** | 0.011 | 0.035 | |
| p-value | *0.559* | ***0.023*** | ***0.028*** | *0.771* | *0.144* | ***0.008*** | *0.591* | *0.054* | |

**Table S8. Pairwise comparison of allometry slopes among genera.**
Allometry models' construction and comparison were calculated using the 'procD.lm' and 'anova' functions in 'geomorph', pairwise comparison was performed using the parse function in the RRPP package on the best model. ANOVA tables of the three tested models (common allometry, different intercepts among genera, and different slopes among genera) are reported, along with model comparison. The last model with different slopes among genera was preferred, and the pairwise analyses of absolute slope distance, angle between slopes and difference in vector lengths for each genus was conducted. Size is the log-transformed Centroid Size. P-values for pairwise differences calculated with 1000 permutations. Significant p-values (p<0.05) are in bold.

**ANOVA tables tested models**

*Common allometry*

| | Df | SS | MS | $R^2$ | F | Z | Pr(>F) |
|---|---|---|---|---|---|---|---|
| size | 1 | 0.864 | 0.864 | 0.302 | 25.474 | 5.086 | **0.001** |
| Residuals | 59 | 2.000 | 0.034 | 0.698 | | | |
| Total | 60 | 2.864 | | | | | |

*Different intercepts genera*

| | Df | SS | MS | $R^2$ | F | Z | Pr(>F) |
|---|---|---|---|---|---|---|---|
| size | 1 | 0.864 | 0.864 | 0.302 | 80.478 | 6.675 | **0.001** |
| genus | 8 | 1.453 | 0.182 | 0.507 | 16.924 | 6.360 | **0.001** |
| Residuals | 51 | 0.547 | 0.011 | 0.191 | | | |
| Total | 60 | 2.864 | | | | | |

*Different slopes genera*

| | Df | SS | MS | $R^2$ | F | Z | Pr(>F) |
|---|---|---|---|---|---|---|---|
| size | 1 | 0.864 | 0.864 | 0.302 | 100.478 | 7.096 | **0.001** |
| genus | 8 | 1.453 | 0.182 | 0.507 | 21.130 | 6.630 | **0.001** |
| size:genus | 8 | 0.178 | 0.022 | 0.062 | 2.584 | 3.334 | **0.001** |
| Residuals | 43 | 0.370 | 0.009 | 0.129 | | | |
| Total | 60 | 2.864 | | | | | |

**ANOVA models' comparison**

| | ResDf | Df | RSS | SS | MS | $R^2$ | F | Z | P |
|---|---|---|---|---|---|---|---|---|---|
| Common allometry (Null) | 59 | 1 | 2.000 | 0.000 | | | | | |
| Different intercepts genera | 51 | 8 | 0.547 | 1.453 | 0.182 | 0.507 | 16.924 | 6.360 | **0.001** |
| Different slopes genera | 43 | 16 | 0.370 | 1.631 | 0.102 | 0.569 | 11.857 | 7.826 | **0.001** |
| Total | 60 | 2.864 | | | | | | | |

**Pairwise comparisons genera by stage**

| | *Absolute distances between slopes* | | | *Angular differences between slopes* | | | *Differences in slope vector length* | | |
|---|---|---|---|---|---|---|---|---|---|
| | d | Z | Pr>d | angle | Z | Pr>angle | d | Z | Pr>d |
| *Balaena:Balaenoptera* (l.) | 0.143 | 0.258 | 0.402 | 36.315 | 0.598 | 0.280 | 0.036 | 0.109 | 0.467 |
| *Balaena:Balaenoptera* (s.) | 0.189 | 0.742 | 0.220 | 42.830 | 0.667 | 0.249 | 0.031 | 0.056 | 0.498 |
| *Balaena:Caperea* | 0.331 | -0.770 | 0.778 | 53.145 | -0.968 | 0.818 | 0.174 | -0.006 | 0.497 |
| *Balaena:Eschrichtius* | 0.315 | 0.180 | 0.438 | 47.317 | -0.928 | 0.818 | 0.183 | 0.914 | 0.187 |
| *Balaena:Kogia* | 0.350 | 2.887 | **0.002** | 59.962 | 2.061 | **0.020** | 0.162 | 2.301 | **0.005** |
| *Balaena:Lagenorhynchus* | 0.303 | 1.227 | 0.111 | 57.152 | 0.823 | 0.207 | 0.116 | 1.200 | 0.121 |
| *Balaena:Megaptera* | 0.138 | -0.140 | 0.567 | 35.117 | 0.215 | 0.424 | 0.043 | 0.359 | 0.366 |
| *Balaena:Phocoena* | 0.314 | 1.592 | 0.059 | 59.484 | 1.071 | 0.150 | 0.118 | 1.405 | 0.087 |
| *Balaenoptera* (l.):*Balaenoptera* (s.) | 0.120 | 1.113 | 0.143 | 24.289 | 0.205 | 0.424 | 0.067 | 1.816 | **0.026** |
| *Balaenoptera* (l.):*Caperea* | 0.350 | -0.226 | 0.598 | 57.708 | -0.138 | 0.544 | 0.210 | 0.030 | 0.500 |
| *Balaenoptera* (l.):*Eschrichtius* | 0.334 | 0.761 | 0.227 | 50.725 | 0.035 | 0.475 | 0.219 | 0.977 | 0.157 |
| *Balaenoptera* (l.):*Kogia* | 0.313 | 3.956 | **0.001** | 49.963 | 2.975 | **0.001** | 0.198 | 3.967 | **0.001** |
| *Balaenoptera* (l.):*Lagenorhynchus* | 0.285 | 1.595 | 0.053 | 52.965 | 1.259 | 0.108 | 0.152 | 1.413 | 0.073 |
| *Balaenoptera* (l.):*Megaptera* | 0.071 | 0.099 | 0.470 | 20.276 | 0.703 | 0.232 | 0.007 | -0.655 | 0.751 |
| *Balaenoptera* (l.):*Phocoena* | 0.260 | 1.653 | 0.050 | 45.626 | 0.776 | 0.224 | 0.154 | 1.732 | **0.032** |
| *Balaenoptera* (s.):*Caperea* | 0.389 | 0.076 | 0.476 | 65.312 | 0.272 | 0.376 | 0.143 | -0.522 | 0.711 |

| | | | | | | | | | |
|---|---|---|---|---|---|---|---|---|---|
| *Balaenoptera* (s.):*Eschrichtius* | 0.377 | 1.031 | 0.137 | 61.135 | 0.619 | 0.281 | 0.152 | 0.473 | 0.310 |
| *Balaenoptera* (s.):*Kogia* | 0.344 | 4.594 | 0.001 | 57.488 | 3.012 | 0.001 | 0.131 | 2.792 | 0.001 |
| *Balaenoptera* (s.):*Lagenorhynchus* | 0.294 | 1.522 | 0.060 | 53.889 | 1.023 | 0.151 | 0.085 | 0.710 | 0.247 |
| *Balaenoptera* (s.):*Megaptera* | 0.135 | 1.135 | 0.135 | 28.182 | 0.343 | 0.366 | 0.074 | 1.925 | 0.017 |
| *Balaenoptera* (s.):*Phocoena* | 0.273 | 1.569 | 0.054 | 49.057 | 0.689 | 0.253 | 0.086 | 0.897 | 0.200 |
| *Caperea:Eschrichtius* | 0.393 | -0.800 | 0.784 | 56.016 | -1.362 | 0.908 | 0.009 | -1.709 | 0.954 |
| *Caperea:Kogia* | 0.469 | 0.762 | 0.215 | 70.210 | 0.611 | 0.267 | 0.012 | -3.151 | 0.997 |
| *Caperea:Lagenorhynchus* | 0.446 | 0.281 | 0.382 | 70.305 | 0.243 | 0.396 | 0.058 | -0.982 | 0.838 |
| *Caperea:Megaptera* | 0.342 | -0.366 | 0.648 | 55.096 | -0.426 | 0.655 | 0.217 | 0.131 | 0.446 |
| *Caperea:Phocoena* | 0.452 | 0.371 | 0.352 | 71.269 | 0.326 | 0.375 | 0.056 | -1.070 | 0.866 |
| *Eschrichtius:Kogia* | 0.435 | 1.485 | 0.073 | 63.608 | 0.801 | 0.199 | 0.021 | -1.623 | 0.949 |
| *Eschrichtius:Lagenorhynchus* | 0.438 | 1.118 | 0.130 | 67.616 | 0.560 | 0.281 | 0.067 | -0.100 | 0.546 |
| *Eschrichtius:Megaptera* | 0.343 | 0.801 | 0.222 | 52.929 | 0.126 | 0.447 | 0.226 | 1.064 | 0.135 |
| *Eschrichtius:Phocoena* | 0.429 | 1.091 | 0.133 | 65.957 | 0.440 | 0.339 | 0.066 | -0.095 | 0.546 |
| *Kogia:Lagenorhynchus* | 0.316 | 1.712 | **0.049** | 48.802 | 0.581 | 0.283 | 0.046 | -0.038 | 0.519 |
| *Kogia:Megaptera* | 0.310 | 3.867 | **0.001** | 48.832 | 2.631 | **0.005** | 0.205 | 4.006 | **0.001** |
| *Kogia:Phocoena* | 0.304 | 1.965 | **0.025** | 46.646 | 0.460 | 0.322 | 0.044 | -0.004 | 0.523 |
| *Lagenorhynchus:Megaptera* | 0.275 | 1.425 | 0.079 | 50.156 | 0.906 | 0.190 | 0.159 | 1.521 | 0.061 |
| *Lagenorhynchus:Phocoena* | 0.250 | 0.010 | 0.494 | 40.903 | -1.242 | 0.887 | 0.001 | -2.088 | 0.978 |
| *Megaptera:Phocoena* | 0.275 | 1.772 | **0.041** | 49.810 | 1.006 | 0.148 | 0.160 | 1.845 | **0.020** |

**Table S9. Pairwise comparison of allometry slopes in ancestral nodes and genera.**
A regression model was calculated with the 'lm' function in base R recreating the best model found for extant taxa in the allometry analysis (formula = RegScores ~ logCS * genus/node). The regression scores and logCS for extant taxa were extracted from the allometry model of size with slopes varying among genera. Regression scores for the ancestral nodes were calculated based on the ancestral intercepts and slopes estimate with the 'fastAnc' function in the package 'phytools' using a distribution of logCS values based on the range of modern taxa. Each node was considered as a group to be compared with the extant genera. Nodes are numbered as follows: 0 – Cetacea, 1 – Mysticeti, 2 – *Caperea*, 3 – Balaenopteridae, 4 – Eschrichtiidae, 5 – *Megaptera*, 6 – Odontoceti, 7 – Delphinoidea. Pairwise comparison was performed in the package emmeans. Upper triangle: Tukey-adjusted P-values of mean comparisons, Diagonal: emmeans estimates for each group (italicized), Lower triangle: comparisons of emmeans estimates earlier vs. later. Pairwise comparisons among extant genera are shaded as the values obtained using the pairwise function on the original 'procD.lm' model are more reliable than these emmeans values. Significant p-values (p<0.05) are in bold.

| | 0 | 1 | 2 | 3 | 4 | 5 | 6 | 7 | Bala | Bal (l.) | Bal (s.) | Cap | Esc | Kog | Lag | Meg | Pho |
|---|---|---|---|---|---|---|---|---|---|---|---|---|---|---|---|---|---|
| 0 | *[0.017]* | **<.0001** | **<.0001** | **<.0001** | **<.0001** | **<.0001** | **0.044** | **<.0001** | **<.0001** | **<.0001** | 0.916 | 0.996 | 0.105 | **<.0001** | **<.0001** | 0.485 | 0.992 |
| 1 | -0.016 | *[0.033]* | 0.980 | 1.000 | 0.961 | 0.395 | **<.0001** | **<.0001** | **<.0001** | 0.979 | **<.0001** | 1.000 | **0.001** | **<.0001** | **<.0001** | 0.029 | 1.000 |
| 2 | -0.015 | 0.002 | *[0.031]* | 0.813 | 0.097 | **0.003** | **<.0001** | **<.0001** | **<.0001** | 1.000 | **<.0001** | 1.000 | **0.001** | **<.0001** | **<.0001** | 0.192 | 1.000 |
| 3 | -0.017 | -0.001 | -0.002 | *[0.034]* | 0.999 | 0.760 | **<.0001** | **<.0001** | **<.0001** | 0.900 | **<.0001** | 1.000 | **0.0004** | **<.0001** | **<.0001** | **0.014** | 0.999 |
| 4 | -0.018 | -0.002 | -0.004 | -0.001 | *[0.035]* | 1.000 | **<.0001** | **<.0001** | **<.0001** | 0.408 | **<.0001** | 1.000 | **0.0003** | **<.0001** | **0.0002** | **0.002** | 0.996 |
| 5 | -0.019 | -0.003 | -0.005 | -0.002 | -0.001 | *[0.036]* | **<.0001** | **<.0001** | **<.0001** | 0.103 | **<.0001** | 1.000 | **0.0002** | **<.0001** | **0.0004** | **0.0003** | 0.986 |
| 6 | 0.004 | 0.020 | 0.018 | 0.021 | 0.022 | 0.023 | *[0.013]* | **<.0001** | **<.0001** | **<.0001** | 1.000 | 0.923 | 0.242 | **<.0001** | **<.0001** | **0.009** | 0.836 |
| 7 | 0.011 | 0.027 | 0.026 | 0.028 | 0.029 | 0.030 | 0.007 | *[0.006]* | **<.0001** | **<.0001** | **0.042** | 0.347 | 0.679 | **<.0001** | **<.0001** | **<.0001** | 0.137 |
| Bala | 0.066 | 0.050 | 0.052 | 0.050 | 0.048 | 0.047 | 0.070 | 0.077 | *[0.083]* | <.0001 | <.0001 | <.0001 | <.0001 | <.0001 | 0.494 | <.0001 | <.0001 |
| Bal (l.) | 0.014 | -0.003 | -0.001 | -0.003 | -0.005 | -0.006 | 0.017 | 0.025 | 0.053 | *[0.030]* | <.0001 | 1.000 | 0.002 | <.0001 | <.0001 | 0.651 | 1.000 |
| Bal (s.) | -0.004 | -0.020 | -0.018 | -0.021 | -0.022 | -0.023 | 0.000 | 0.007 | 0.070 | 0.017 | *[0.013]* | 0.938 | 0.256 | <.0001 | <.0001 | 0.083 | 0.875 |
| Cap | 0.011 | -0.005 | -0.004 | -0.006 | -0.007 | -0.008 | 0.015 | 0.022 | 0.056 | 0.003 | -0.015 | *[0.028]* | 0.058 | <.0001 | 0.038 | 1.000 | 1.000 |
| Esc | -0.037 | -0.053 | -0.052 | -0.054 | -0.055 | -0.056 | -0.033 | -0.026 | 0.103 | 0.051 | 0.033 | 0.048 | *[-0.020]* | 1.000 | <.0001 | 0.022 | 0.043 |
| Kog | -0.041 | -0.058 | -0.056 | -0.058 | -0.059 | -0.060 | -0.037 | -0.030 | 0.108 | 0.055 | 0.038 | 0.052 | 0.004 | *[-0.024]* | <.0001 | <.0001 | <.0001 |
| Lag | 0.045 | 0.029 | 0.031 | 0.029 | 0.027 | 0.026 | 0.049 | 0.056 | 0.021 | -0.032 | -0.049 | -0.035 | -0.082 | -0.087 | *[0.062]* | <.0001 | 0.003 |
| Meg | 0.007 | -0.010 | -0.008 | -0.010 | -0.011 | -0.013 | 0.011 | 0.018 | 0.060 | 0.007 | -0.010 | 0.004 | -0.044 | -0.048 | 0.039 | *[0.024]* | 1.000 |
| Pho | 0.009 | -0.007 | -0.005 | -0.007 | -0.009 | -0.010 | 0.013 | 0.020 | 0.057 | 0.004 | -0.013 | 0.002 | -0.046 | -0.051 | 0.036 | -0.003 | *[0.026]* |

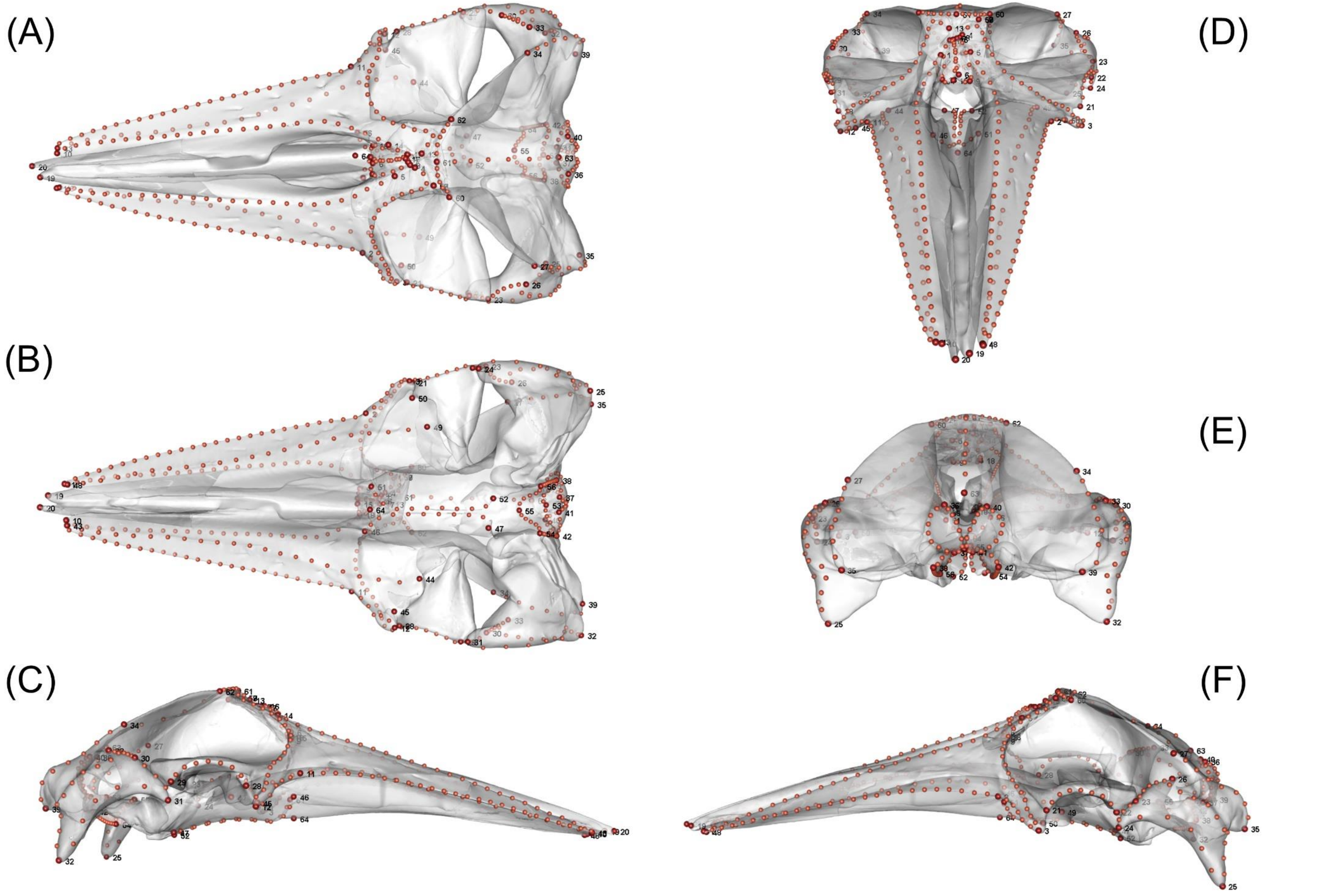


**Figure S1.** Landmarking scheme used in this study, plotted on the skull of an adult *Balaenoptera* (*B. acutorostrata*, CAS 23867). (A) Dorsal view; (B) Ventral view; (C) Medial view; (D) Anterior view; (E) Posterior view; (F) Lateral view. Fixed landmark points in red and numbered, curve semi-landmarks in orange. See Table S3 for details.

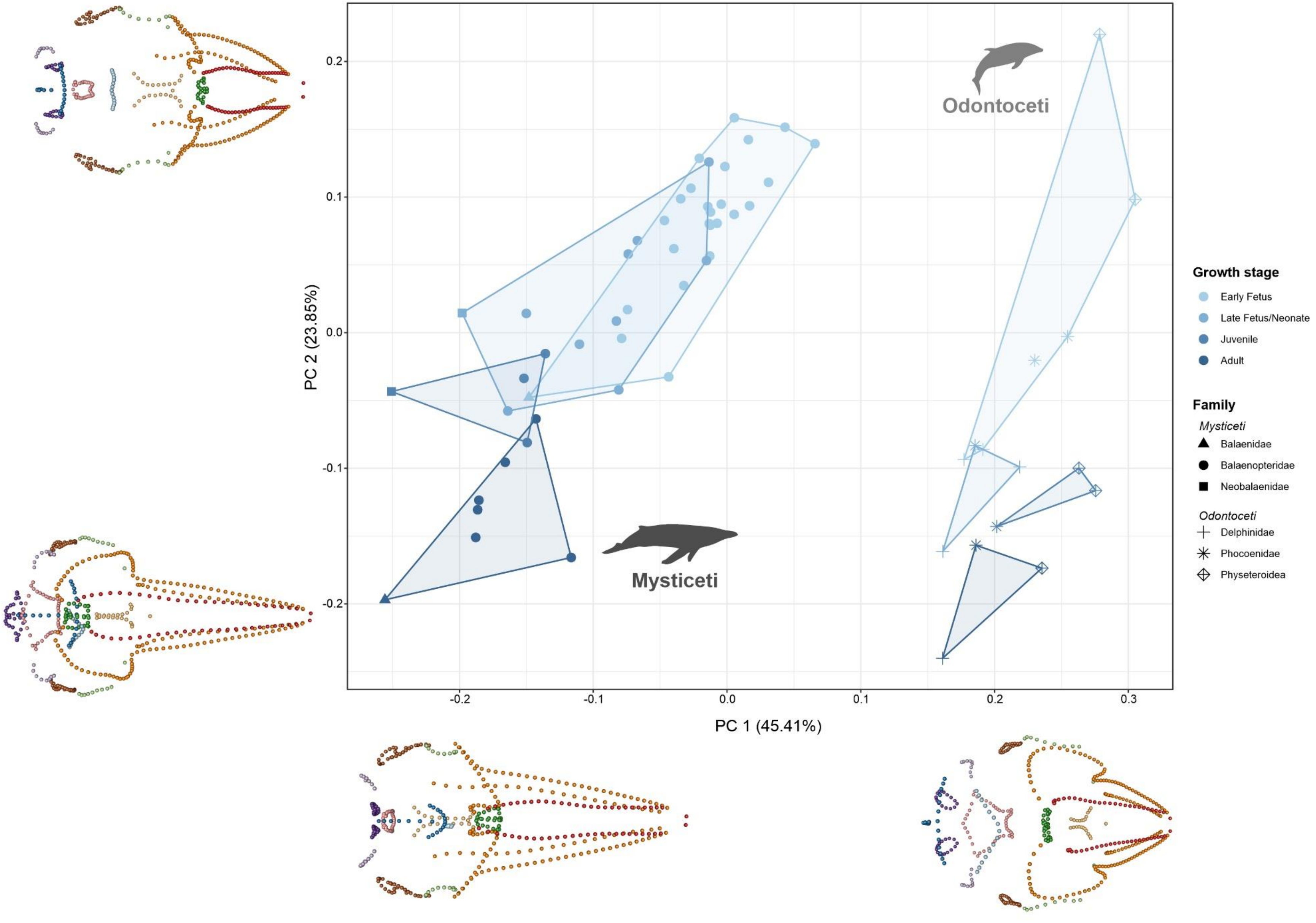


**Figure S2**. Skull ontogeny morphospace (PCA), showing the shape variation through ontogeny and phylogeny in the full raw dataset. Points and areas are coloured based on ontogenetic stage in each group, shapes represent families. The landmark plots on the left and on the bottom represent the morphological extremes on the PC1 and PC2 axis, in dorsal view. Fixed landmarks and semi-landmarks are represented in unique colours to highlight shape changes in different regions of the skull.

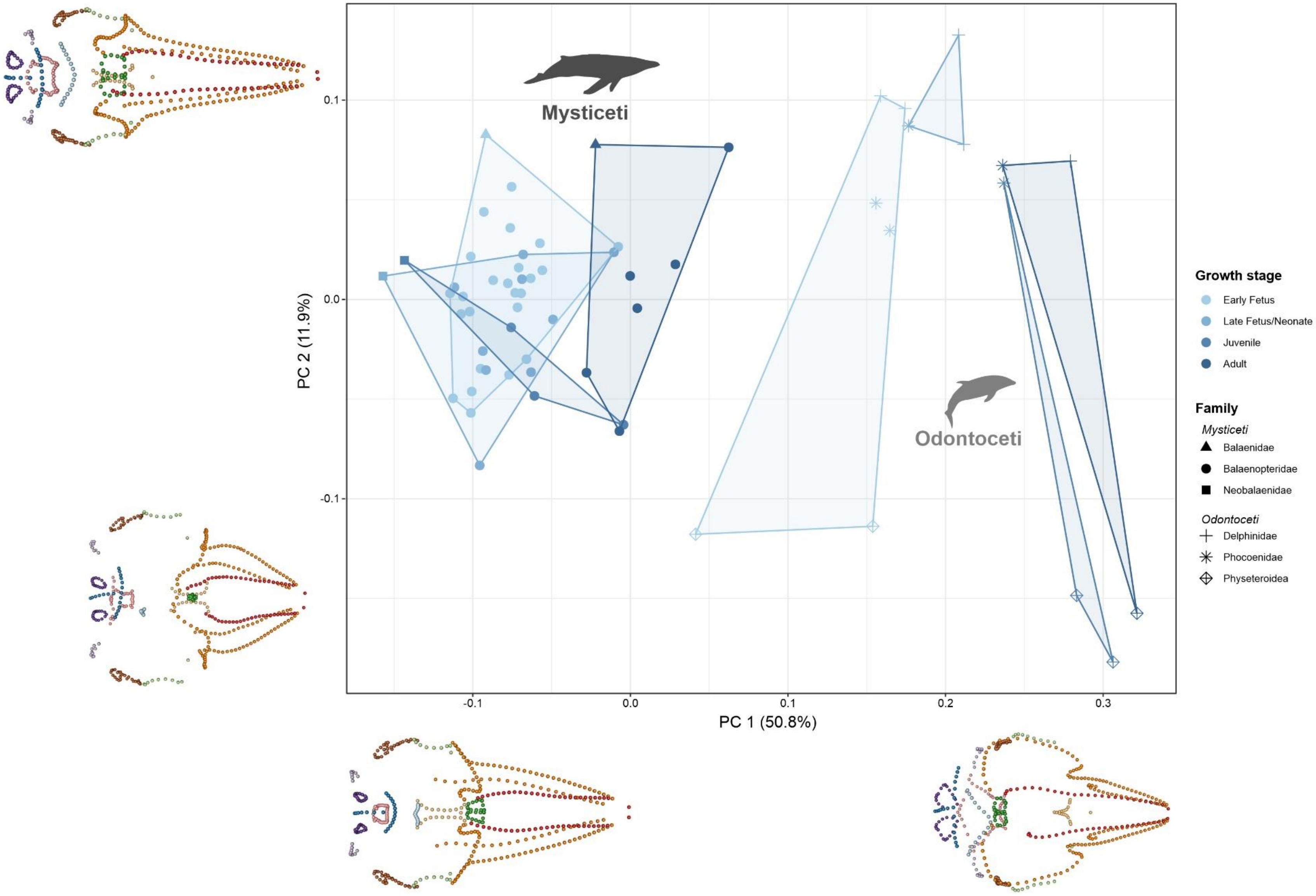


**Figure S3**. Skull ontogeny morphospace (PCA), showing the shape variation through ontogeny and phylogeny in the full allometry residuals dataset. Points and areas are coloured based on ontogenetic stage in each group, shapes represent families. The landmark plots on the left and on the bottom represent the morphological extremes on the PC1 and PC2 axis, in dorsal view. Fixed landmarks and semi-landmarks are represented in unique colours to highlight shape changes in different regions of the skull, see Figure 1 for details.

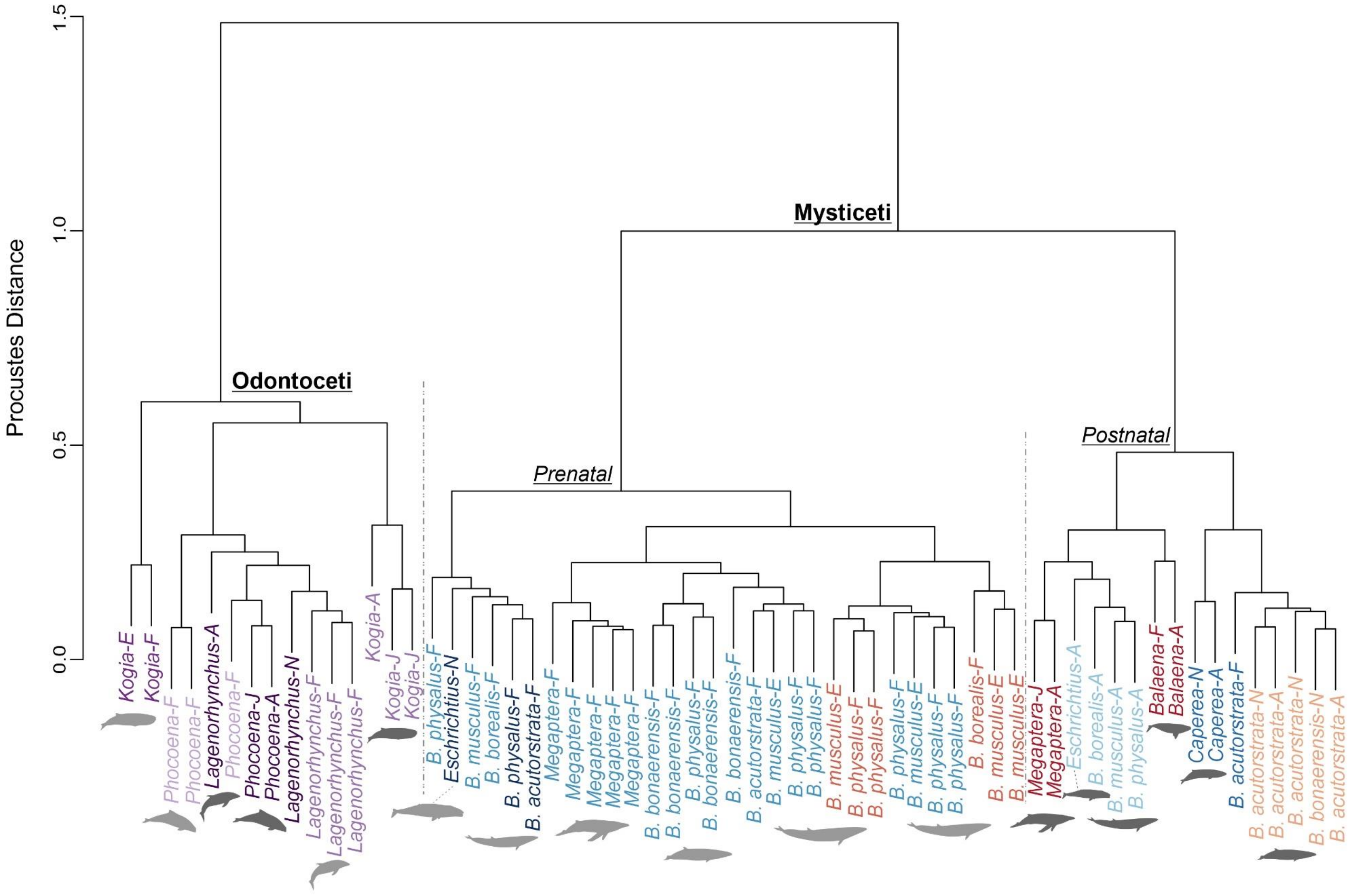


**Figure S4**. Clustering analyses on full dataset. Dendrogram of Procrustes distances obtained from 'hclust' function in the 'stats' package using Ward's method. Tip labels coloured according to the results of the k-mean clustering on shape data performed with the 'LoydShapes' function in the 'Anthropometry' package with k=10. The main clusters identifiable in the dendrogram are labelled. Specimens are labelled with the species names and their ontogenetic age (E = embryo, F = fetus, N = neonate, J = juvenile, A = adult). The silhouettes indicate the position of major clusters of each taxon (light grey = prenatal cluster, dark grey = postnatal cluster).

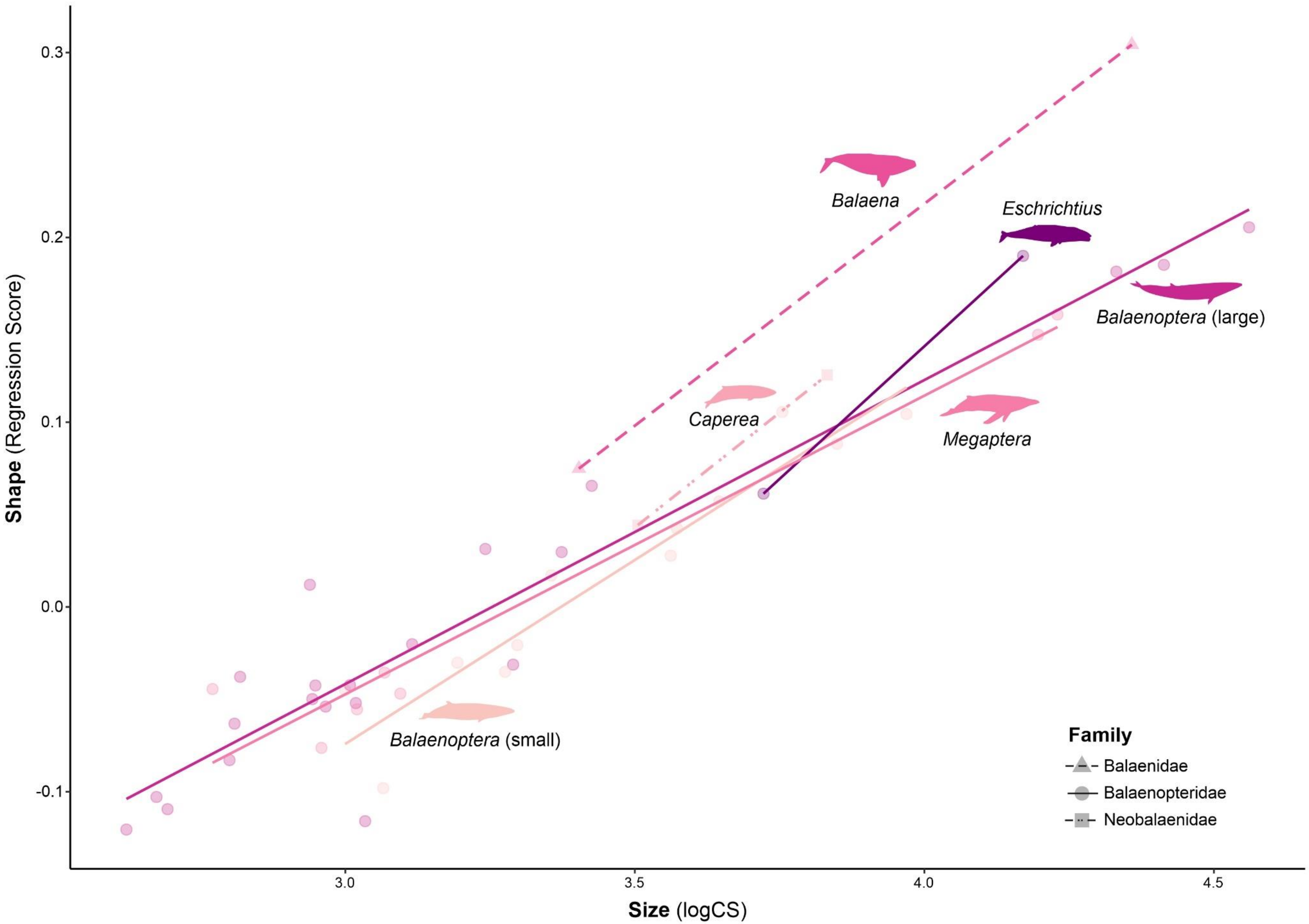


**Figure S5.** Allometric regression of skull shape and size (logCS) by genus for Mysticeti. Model was calculated using the 'procD.lm' function in 'geomorph' (formula = RegScores ~ logCS * genus). Families are represented by different shapes and line types. See Table S8 for details on the model and pairwise comparison of slopes.

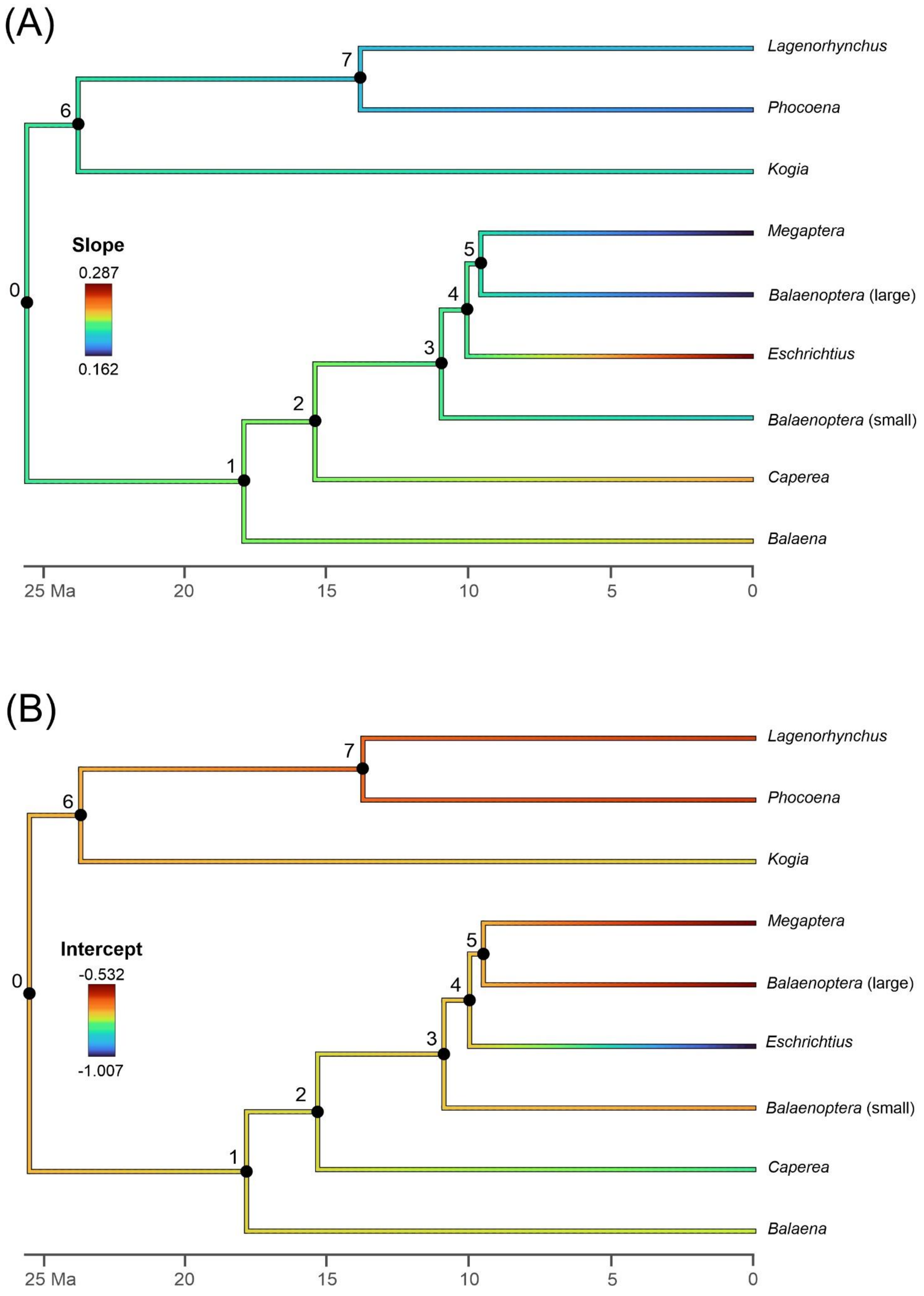


**Figure S6.** Ancestral state reconstruction of allometry (A) slope and (B) intercept values in the full dataset. Ancestral states at the nodes were estimated with the 'fastAnc' function in 'phytools' starting from the data extracted from the 'procD.lm' model (see Figure S5, Table S8 for details). Character changes were mapped on the tree with the 'contMap' function in 'phytools'. Tree topology and node ages from McGowen et al. (2020).

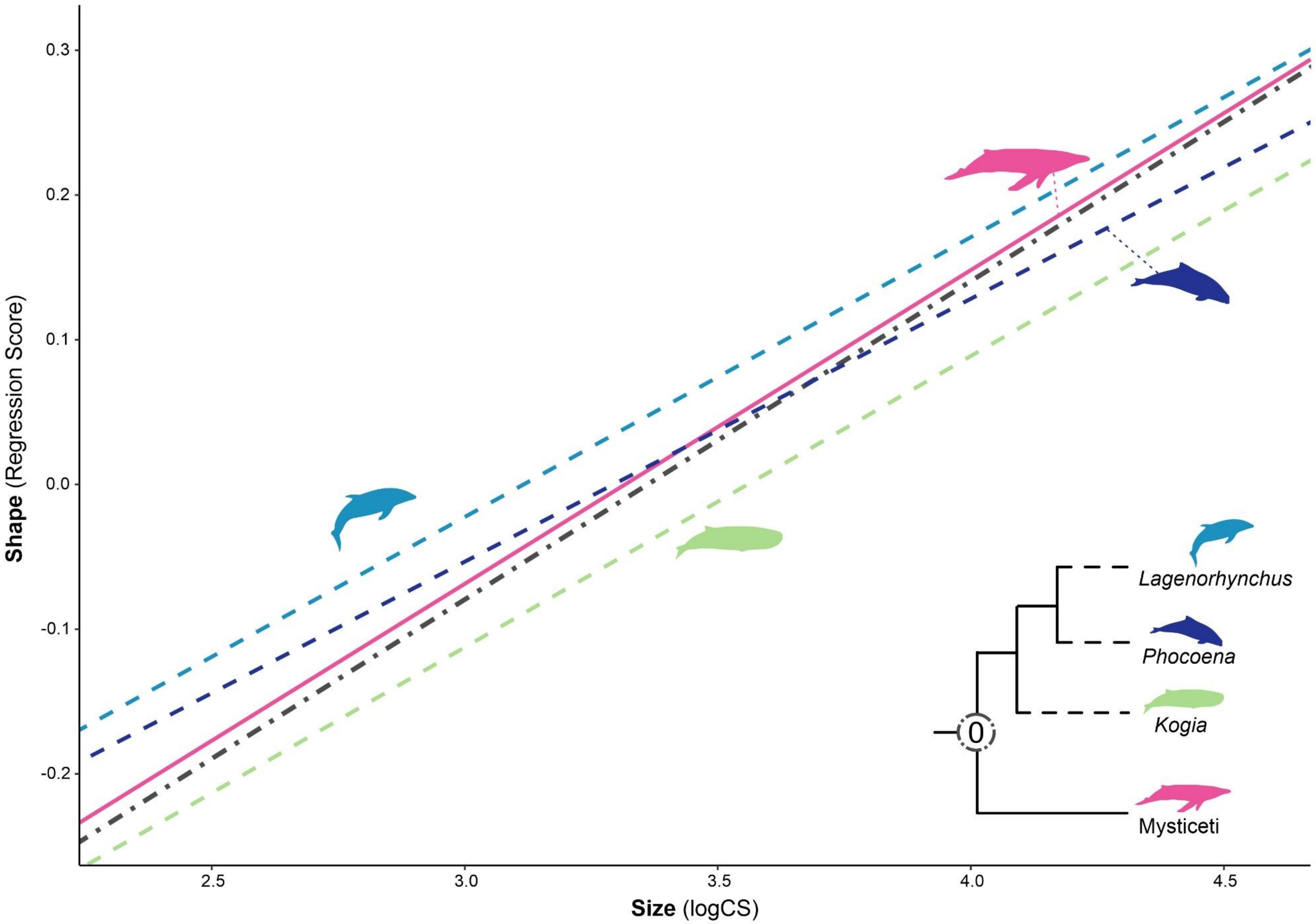


**Figure S7.** Allometric regression of skull shape and size (logCS) of all taxa and their common ancestor. Model for modern taxa calculated using the 'procD.lm' function in 'geomorph' (formula = RegScores ~ logCS * genus). Ancestral allometry regression parameters for all Cetacea (node 0) were calculated using 'fastAnc' function in 'phytools'. The two groups and node are represented by different colors and line types. Mysticeti is plotted as the average regression for all taxa to allow for an easier comparison with the odontocetes. See Table S9 for pairwise comparison of slopes.

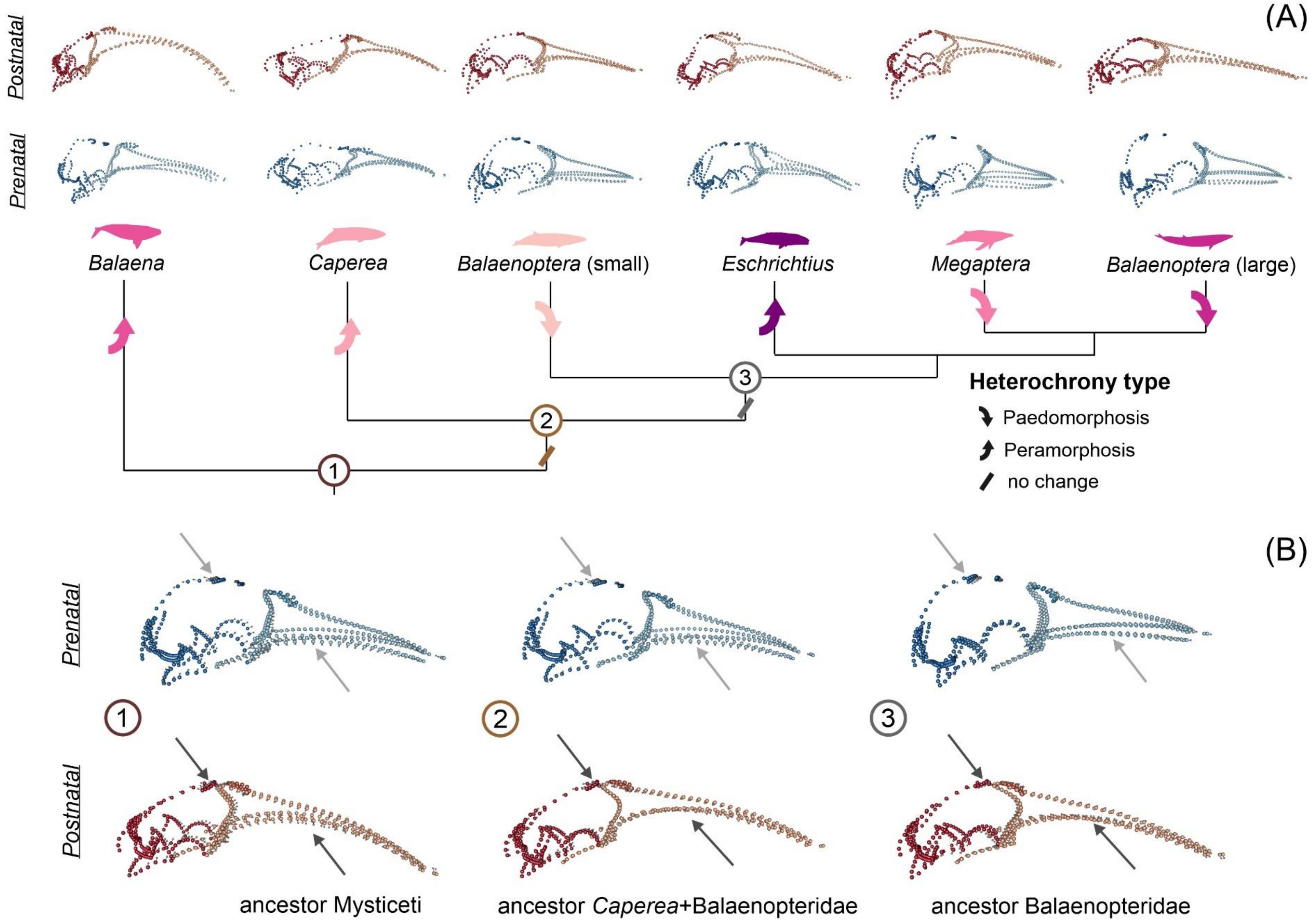


**Figure S8.** Heterochrony in Mysticeti phylogeny and related rostrum arching changes. (A) Phylogeny with inferred direction of heterochronic change for each node and mean shapes for prenatal (blue) and postnatal (red) stages of each taxon; (B) reconstructed prenatal and postnatal skull shapes at the ancestral nodes. Landmarks in the neurocranial region are highlighted with a darker shade, nasals have a medium shade, and the rostrum has a lighter shade. Shapes for extant taxa are estimated mean shapes for all prenatal and postnatal specimens for each taxon, ancestral shapes for qualitive comparison were reconstructed by conducting separate phyloPCA analyses for prenatal and postnatal stages using the relative mean shapes. The arrows on the ancestral skull shapes point to the region where the developmental differences with modern taxa are more marked, as described in the text. Skull shapes in medial view (dorsal view in Figure 4), not to scale.
